# RNA Infrastructure Profiling Illuminates Transcriptome Structure in Crowded Spaces

**DOI:** 10.1101/2023.10.09.561413

**Authors:** Lu Xiao, Linglan Fang, Eric T. Kool

## Abstract

RNAs can fold into compact three-dimensional structures, and most RNAs undergo protein interactions in the cell. These compact and occluded environments can block the ability of structure-probing agents to provide useful data about the folding and modification of the underlying RNA. The development of probes that can analyze structure in crowded settings, and differentiate the proximity of interactions, can shed new light on RNA biology. To this end, here we employ 2,-OH-reactive probes that are small enough to access folded RNA structure underlying many close molecular contacts within cells, providing considerably broader coverage for intracellular RNA structural analysis. We compare reverse transcriptase stops in RNA-Seq data from probes of small and standard size to assess RNA-protein proximity and evaluate solvent-exposed tunnels adjacent to RNA. The data are analyzed first with structurally characterized complexes (human 18S and 28S RNA), and then applied transcriptome-wide to polyadenylated transcripts in HEK293 cells. In our transcriptome profile, the smallest probe acetylimidazole (AcIm) yields 80% greater structural coverage than larger conventional reagent NAIN3, providing enhanced structural information in hundreds of transcripts. We further show that acetyl probes provide superior signals for identifying m^6^A modification sites in transcripts, and provide information regarding methylation sites that are inaccessible to a larger standard probe. RNA infrastructure profiling (RISP) enables enhanced analysis of transcriptome structure, modification, and interactions in living cells, especially in spatially crowded settings.

## Introduction

RNA plays a critical role in many cellular processes, ranging from gene translation to regulation, both in normal and pathological conditions (1, 2). The diversity of RNA structures, from simple stem-loop configurations to intricate 3D structures and complex assemblies with proteins, empowers RNAs to modulate cell functions across varied conditions and cell types (3). Understanding the relationship between RNA structure and function is an essential part of decoding transcriptome biology, and is also important for targeting RNAs for disease treatment (4-6). To shed light on RNA structure, chemical probing techniques have become a widely utilized strategy. This approach takes advantage of reactions of electrophilic probing molecules with nucleophilic groups in RNA. Commonly employed methods include dimethyl sulfate (DMS) footprinting (7), N3-kethoxal probing with deep sequencing (Keth-seq) (8), and selective 2’-hydroxyl acylation analyzed by primer extension (SHAPE) (9). These techniques provide valuable insights into spatial 2D and 3D structures and conformational changes of RNA molecules, aiding in the study of their functional roles and their potential for therapeutic targeting.

Among these approaches, SHAPE methodology, initially developed by Weeks, leverages the elevated acylation reactivity of 2’-hydroxyl (2’-OH) groups in unpaired regions as opposed to paired ones. The acylation adducts of common SHAPE reagents on RNA are analyzed by reverse transcriptase (RT), resulting in stops or mutations that can be detected through deep sequencing (10-12). Various SHAPE reagents with distinct half-lives, cell-permeabilities, and enrichment handles have been developed to enable in-depth and in-cell probing, including two major classes of reagent structures: isatoic anhydrides (e.g., NMIA and 1M7) and acylimidazole reagents (e.g., FAI, NAI, 2A3, NAIN3) (13-17). These reagents have been extensively employed to study secondary/tertiary structures across diverse RNA types and species. In addition to proving data about RNA alone, SHAPE methodology can also provide useful information about RNA-protein interaction by comparison of in-cell profiles with protein-free *in vitro* profiles (16, 18). Importantly, this comparison indicates that the in-cell RNA structural information is often shielded by cellular proteins. Previous studies have shown that ∽95% of total mRNAs possess identified protein-binding sites, and each RNA-binding protein can bind to 5%-30% of over 20,000 transcripts (19). This widespread occlusion by proteins leads to major gaps in data on RNA intracellular structure.

Much of this missing RNA structural data can be attributed to protein contacts, but gaps in data also may arise from other sources such as RNA-RNA folding or partial fragmentation during RNA metabolism. Moreover, it is becoming increasingly evident that many cellular RNAs exist in spatially crowded compartments such as nucleoli, paraspeckles, and stress granules (20, 21). While some information can be gained by comparison of in-cell data with the reconstituted RNA *in vitro*(16), the latter may not always be identical to the structure *in vivo*, as it is surrounded by numerous proteins and other components in cells. For these reasons, the development of tools that can directly probe the infrastructure of RNA underlying close contacts (*e*.*g*., with proteins), can broaden our insights into RNA structures and functions in living cells.

The acylation reactivity of RNA 2’-OH groups is influenced both by the local nucleotide accessibility and by the steric bulk of probing reagents (22, 23). Commonly employed SHAPE reagents contain aromatic rings and other substituents, potentially limiting their access to crowded RNA structure. To address this issue, we hypothesized that probes with small enough size might offer an advantage over standard probes in spatially crowded settings. Among the smallest possible reagents is acetylimidazole (AcIm), which was recently shown to be cell-permeable and was shown to be able to map intracellular rRNA structure (24). In addition, Stephenson *et al*. demonstrated that AcIm is useful as a robust SHAPE reagent capable of generating adducts suitable for nanopore sequencing to achieve single-molecule RNA structural profiles *in vitro* (25). These precedents suggest the potential of this small reagent in probing RNA structures in environments that restrict access to standard probes.

In this work, we test the ability of very small 2’-OH acylating reagents with comparison to a standard-sized agent to probe RNA structures in close macromolecular contacts within cells. This <u>R</u>NA Infra<u>s</u>tructure <u>P</u>rofiling (RISP) method provides a significantly broader depth of information for analysis of intracellular RNA structures. We compared the RT-stop profiles obtained from small and standard-sized RNA structure probes (AcIm or Azido-AcIm (AcIN3) vs. NAIN3) and assessed their probing capability in crowded cellular spaces by analyzing RNA-protein proximity and tunnels adjacent to RNAs in well-characterized 18S and 28S human ribosome complexes. We then expanded the RISP application to measure RNA structural profiles of polyadenylated transcripts throughout the transcriptome in human embryonic stem cells (HEK293). The smallest probe, AcIm, provided significantly more comprehensive folding information of RNAs, particularly in crowded regions associated with close contacts. AcIm also provided enhanced signals for base modification sites that are inaccessible to NAIN3 due to its larger size. The data suggest that RNA infrastructure profiling can enable advanced analysis of transcriptome structure, modifications, and interactions in living cells, particularly in spatially crowded environments.

## Results

### RNA infrastructure profiling (RISP) with probes of small and standard size

Standard in-cell SHAPE reagent NAIN3 has molecular weight 228.21 and volume 193.2 Å^3^, and is ca. 10.6 Å (atom-to-atom) in its largest dimension, whereas AcIm is much smaller (MW 110.12, volume 100.8 Å^3^, atom-to-atom length 6.2 Å) (Fig. S1). AcIN3, slightly larger than AcIm, is previously unknown, and was designed to have a potential pulldown handle (for future use) akin to that of NAI-N3. To investigate the efficacy of these differently-sized 2’-OH acylating reagents in probing intracellular RNA structures with close contacts, we conducted acylation-based measurements on 18S and 28S ribosomal RNAs in intact HEK239 cells using both probes. After optimizing the probing conditions to achieve single-hit kinetics for each reagent, we calculated the R-score as RISP signal at each nucleotide to map the RNA structures. Specifically, we used a modified RBRP pipeline (26) to quantify the reverse transcriptase (RT) stops after library preparation and deep sequencing. RISP in HEK293 cells indicated that both reagents exhibited a strong concordance with manual structure-probing gels (Pearson correlation r≥ 0.85) (Fig. S2a-b), demonstrating that both reagents quantitatively and accurately reflect the known structure of 18S ribosomal RNA (Fig. S2). Notably, small-sized reagents exhibited distinct reactivity towards unpaired nucleotides with close spatial contacts (Fig. S2). Thus, we hypothesized that the information obtained from regions where a small probe displayed dominant signals with a surplus R-score≥ 0.2 over NAIN3 might shed light on RNA infrastructure, that is, structure in crowded spaces associated with close contacts (Fig. 1a, S2).

**Figure 1.**
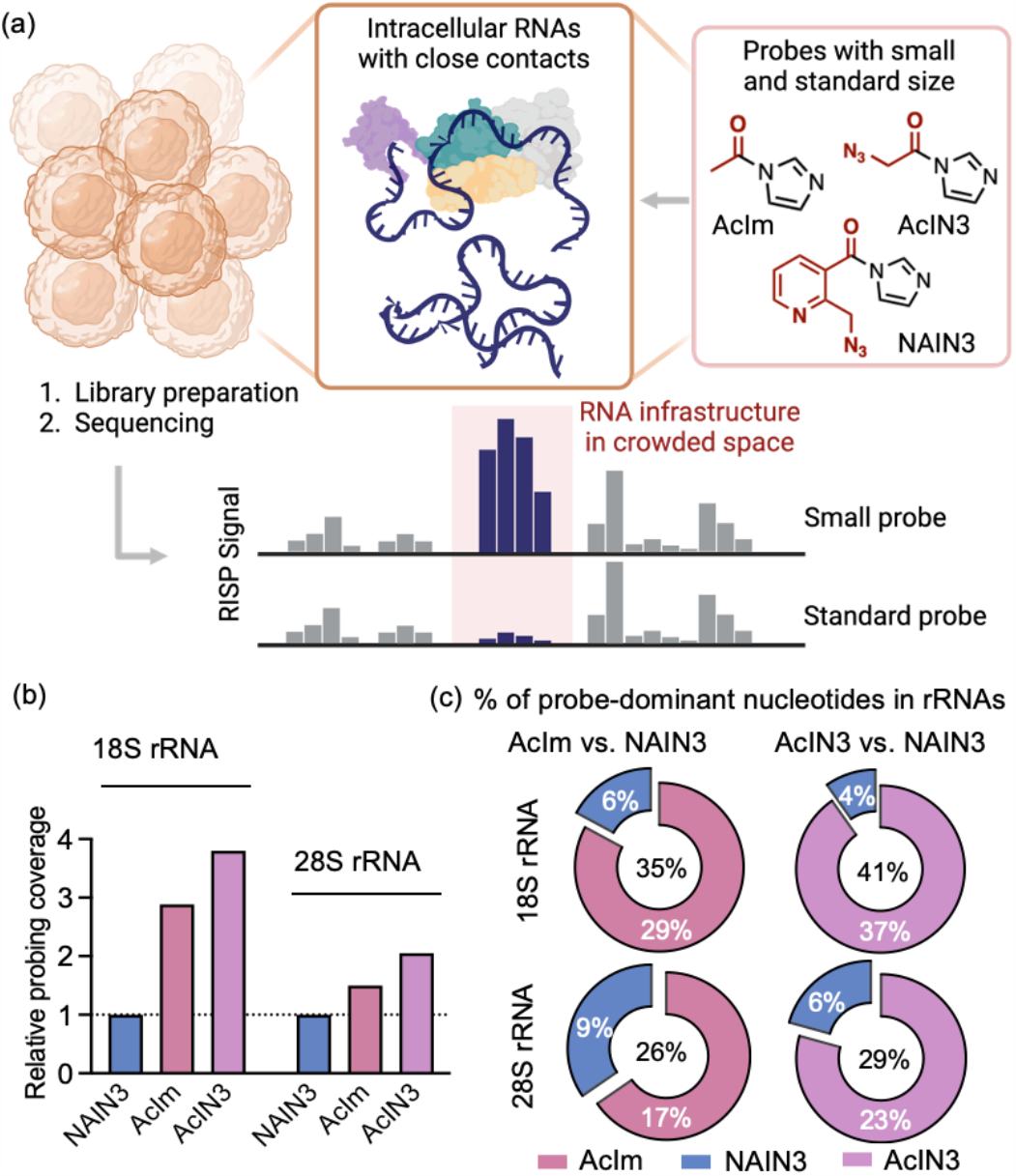
Intracellular RNA infrastructure profiling (RISP) with acylating probes of small and standard size. (a) Schematic of the RISP approach revealing RNA infrastructure in crowded spaces by comparing the sequencing profiles of small (AcIm, AcIN3) and standard (NAIN3) probes. Crowding near 2’-OH groups can arise from close macromolecular contacts, such as RNA-RNA folds and RNA-protein interactions, blocking the access of large probe for acylation. Reagent structures (AcIm, AcIN3, NAIN3) are shown. (b) Overall comparison of relative probing coverage (relative number of nucleotides meeting “efficient” definition) for small (AcIm, AcIN3) and standard (NAIN3) probes in ribosomal RNAs. (c) Percentage of small and standard reagents’ dominant nucleotides in human 18S and 28S rRNAs, showing elevated probing ability, on average, by the small probes. Probe-dominant nucleotides are the ones showing significantly superior signal with one probe having differential R-score larger than 0.2, defined by comparing the calculated R-score with the manual structure-probing gels of partial 18S rRNA for AcIm and NAIN3 reagents.

We first evaluated the relative probing ability of small and standard sized probes (AcIm, AcIN3 and NAIN3) by comparing the number of 2’-OH groups generating efficient signals (R-score≥ 0.1) in 18S and 28S rRNA profiles (Fig. 1b). Both AcIm and AcIN3 exhibited considerably higher probing ability than NAIN3, with ∽3-4 times more signals in 18S rRNA and ∽1.5-2-fold more signals in 28S rRNA, consistent with the highly congested structures of RNA/protein complexation in these macromolecular complexes. Next, we analyzed the percentage of nucleotides in the RNA for which one reagent showed dominant signals in total efficiently probed rRNAs (Fig. 1c). Both acetyl probes, AcIm and AcIN3, displayed a higher proportion of dominant signals in 18S and 28S rRNAs, ranging from 17% to 37%, as compared to NAIN3, which exhibited only 4% to 9% dominant signals. The overall improvement in probing signal with the small acetyl probes prompted us to delve further into exploring the enhanced capabilities of small reagents over larger standard probes.

### Capability of RISP in decoding RNA structures underlying RNA-protein contacts

Having analyzed the dominant signals generated by small and standard probes in rRNAs, we investigated the spatial environment of these dominant nucleotide domains. We speculated that RNA-RNA and RNA-protein interactions might lead to variations in the spatial crowding of RNA assemblies, potentially causing steric hindrance during RNA 2’-OH acylation. Our hypothesis was that due to their smaller sizes, AcIm/AcIN3 dominant signals might primarily arise in spatially crowded regions, whereas NAIN3 dominant nucleotides might be more prevalent in more spacious positions. To test this, we first analyzed published structural data for these macrocomplexes, using the RNA-protein distance of dominant nucleotides to assess the probing signals in the protein interacting space of ribosome. We calculated all distances of rRNA 2’-OH groups to the atoms of nearby proteins in the crystal structure of the human 80S ribosome, using the closest distance as a parameter to evaluate the probe capability (Fig. 2a). Examination of the distributions of the closest distance of small and standard probe-dominant nucleotides (AcIm vs. NAIN3; AcIN3 vs. NAIN3) in rRNAs led to a notable observation: a considerable number of AcIm/AcIN3 dominant nucleotides were found to be located within 5 Å distance to nearby proteins in both 18S and 28S rRNAs, in contrast to much lower frequency of NAIN3 dominant signals in the same range (159 vs. 23 for AcIm vs. NAIN3; 288 vs. 20 for AcIN3 vs. NAIN3; Fig. 2b, S4a). Prior studies have found that many types of RNA-protein interactions, such as hydrophobic interactions, occur within the distance of 5 Å in RNA-protein interfaces (27), which further supports the notion that small-sized reagents may provide improved signals in such contexts.

**Figure 2.**
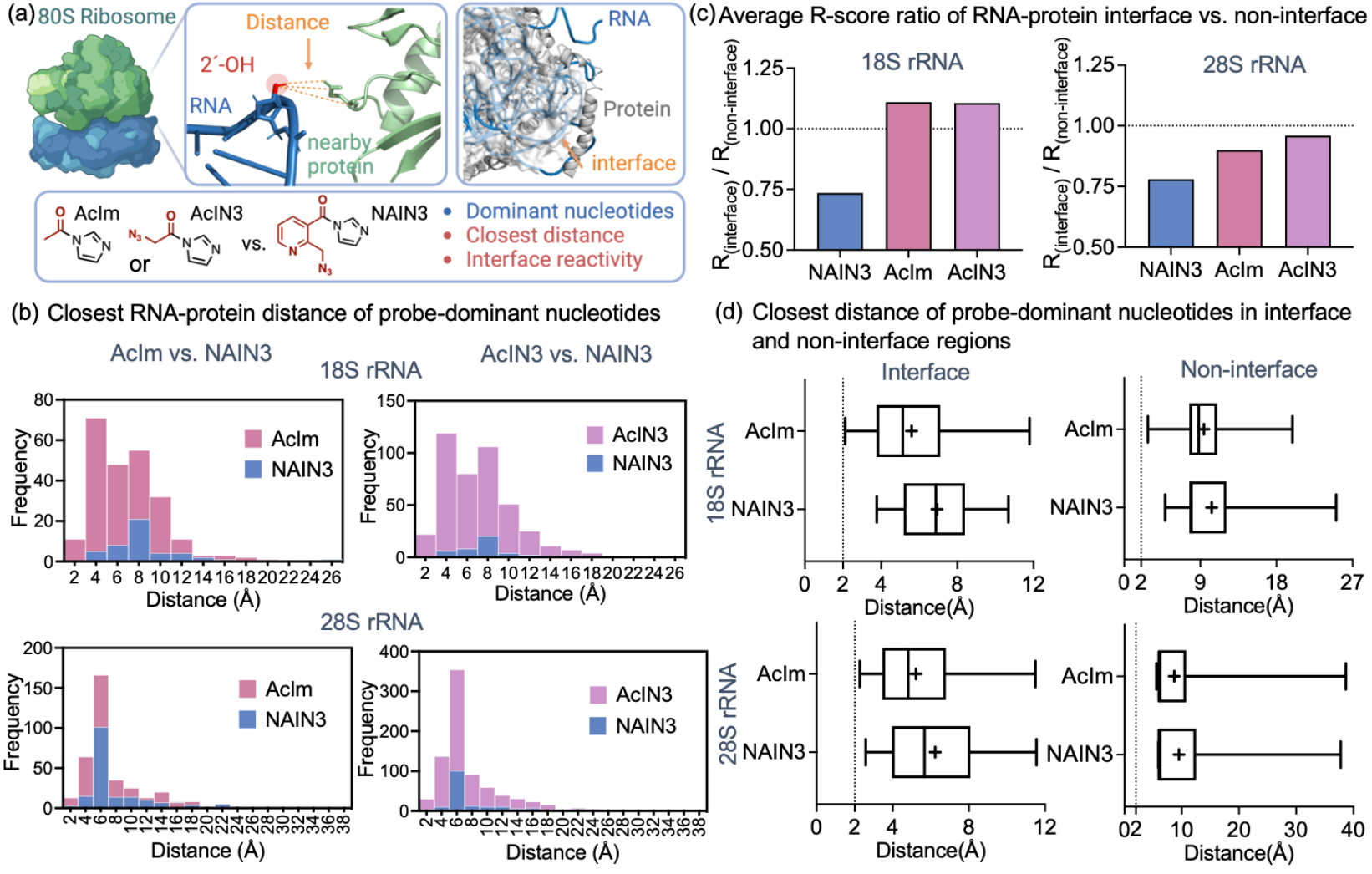
RISP enables structure mapping underlying close RNA-protein contacts. (a) Schematic of probing evaluation for small (AcIm, AcIN3) and standard (NAIN3) reagents in human ribosomal RNAs with associated protein contacts, including RNA-protein distance and RNA-protein interface analyses. (b) Capability evaluation of small (AcIm, AcIN3) and standard (NAIN3) reagents by the closest RNA-protein distance distribution of dominant nucleotides in 18S and 28S rRNAs. (c) Performance evaluation of small (AcIm, AcIN3) and standard (NAIN3) reagents via average reactivity analysis of RNA-protein interfaces versus non-interface RNA regions in human 80S ribosome. (d) Closest RNA-protein distances of probe-dominant nucleotides probed by AcIm and NAIN3 reagents located in the 18S and 28S rRNA-protein interface and non-interface regions.

In addition to measuring distance, we also conducted an RNA-protein interface reactivity analysis of the human ribosome as second parameter to compare the RNA probing ability of small (AcIm, AcIN3) and standard (NAIN3) probes when in contact with proteins (Fig. 2a). We calculated interfaces and non-interfaces residues between RNAs and proteins based on the crystal structure of ribosome. We then compared the reactivity of interface vs. non-interface 2’-OH groups by averaging the nucleotide R-score of AcIm, AcIN3 and NAIN3 in each region (Fig. 2c). The results revealed that AcIm/AcIN3 exhibited higher reactivity in the RNA-protein interfaces for 18S rRNA and similar reactivity for 28S rRNA, as compared to the non-interface regions. In contrast, NAIN3 showed approximately 25% lower reactivity in the interface regions compared to the non-interfaces for both 18S and 28S rRNA complexes. Upon further examination of the closest RNA-protein distance of acetyl or NAIN3 dominant nucleotides in both interface and non-interface regions, we observed a significant difference between acetyl probes and NAIN3 in the interface region of both rRNA complexes (Fig. 2d, S4b). A closest RNA-protein distance as short as ∽2 Å was observed for acetyl dominant nucleotides in distribution, revealing that acetyl probes could efficiently react with nucleotides positioned quite close to proteins. Conversely, in the non-interface regions, the distance distribution of acetyl probes and NAIN3 showed similar or near-identical patterns (Fig. 2d, S4b). Thus the structure/reactivity analyses establish that the small-size acylating reagents have enhanced capability in probing RNA structures amid crowded RNA-protein contacts.

### Evaluation of RISP in association with solvent-exposed tunnels adjacent to RNAs

We wondered whether the smaller size of the acetyl probes could enhance their ability to probe RNA nucleotides situated adjacent to narrow solvent-accessible passageways (tunnels) in the complex 80S ribosome. The 3D structures of internal passageways are defined by spaces between RNA-RNA interactions and RNA-protein interactions (28, 29). These passageways, referred to as solvent-exposed tunnels, exhibit varied dimensional space along their length. One such tunnel, the ribosome exit tunnel, plays an important role in nascent polypeptide folding and translational stalling (30-32). The biological roles of other ribosomal tunnels remain unclear, although they do contribute to solvation of the complex by water and ions. To study this question using probe data, we analyzed the cross-sectional tunnel radius at the positions of small (AcIm, AcIN3) and standard (NAIN3) reagent dominant nucleotides that appear in the tunnels in 18S and 28S rRNA complexes (Fig. 3a). 216 calculated tunnels were examined, including 2068 nucleotides. Among them, ∽40-50% of nucleotides were efficiently probed by AcIm/AcIN3 and NAIN3. AcIm/AcIN3 showed remarkably higher reactivity in tunnel nucleotides compared to NAIN3, with probe-dominant nucleotide percentages of 30.2% (acetyl probes) vs. 2.4% (NAIN3) in 18S rRNA and 17.3% vs. 5.1% in 28S rRNA (Fig. 3b, S5a). The difference was less pronounced in the 28S rRNA complex, likely due to higher crowding effects, consistent with the observation above. We then analyzed the tunnel radius at each of those dominant nucleotides for each reagent (Fig. 3c, S5b). The results revealed that the smaller sizes of AcIm/AcIN3 allowed them to probe nucleotides within narrower tunnels effectively, showing a smaller tunnel radius (down to ∽1 Å) of dominant sites in the distribution. This further underscored the capability of small probes to provide RNA structural information that is otherwise inaccessible to larger standard probes.

**Figure 3.**
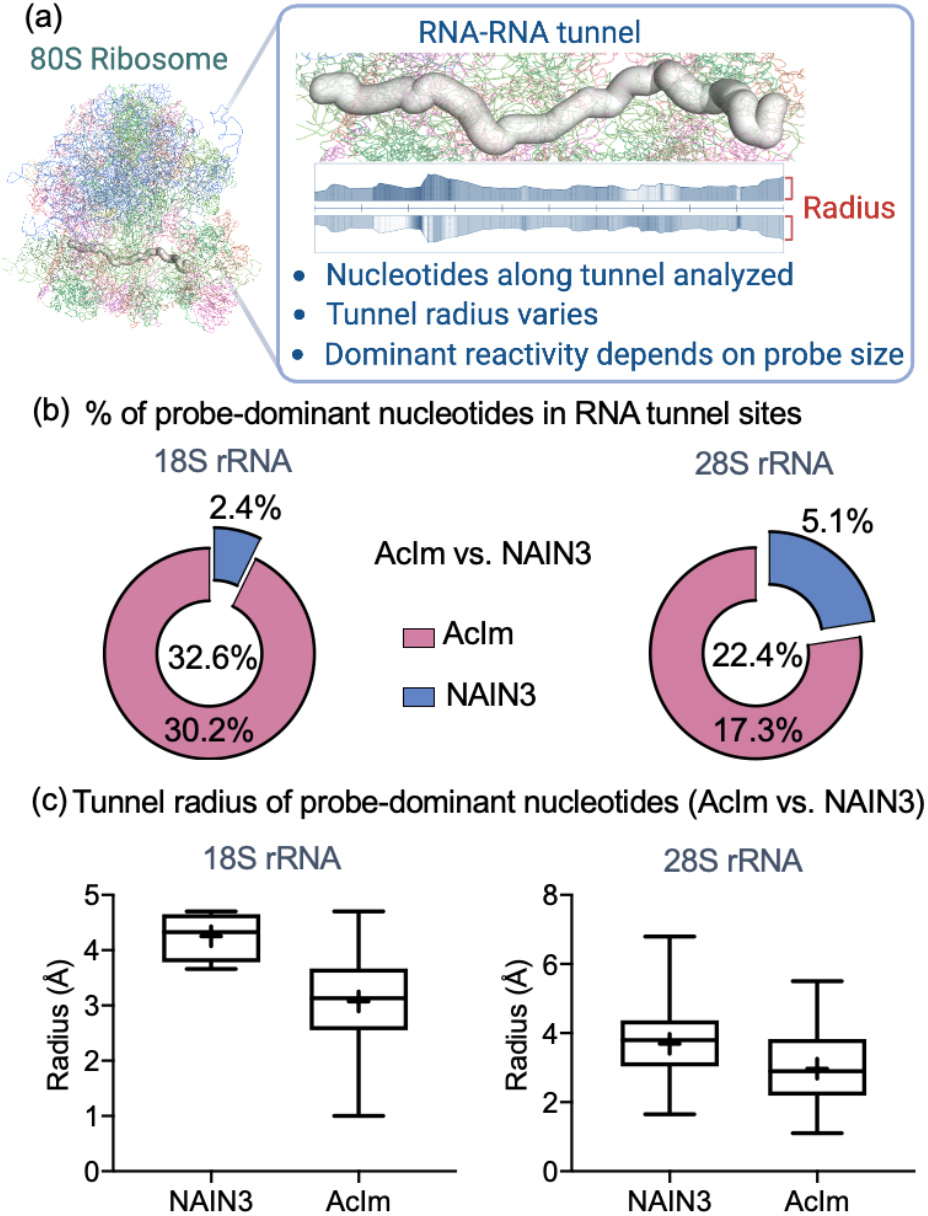
RISP evaluation of probing capability along solvent-exposed tunnels within the ribosomal ribonucleoprotein (RNP) complex. Tunnels arise from RNA-RNA folds and RNA-protein interactions. (a) Schematic of tunnel nucleotides analysis for RISP within the human 80S ribosome. (b) Percentage of small (AcIm) and standard (NAIN3) reagent-dominant nucleotides in probed RNA tunnel sites for 18S and 28S rRNA, showing dominance for small acetyl probes in this tight space; (c) Tunnel radius analysis of AcIm and NAIN3 dominant nucleotides in probed RNA tunnel sites for 18S and 28S rRNA, showing enhanced probing by small probes in tighter tunnel spaces.

### RISP provides enhanced structural information in RNAs transcriptome-wide

The above data established the capability of RISP with small acylating reagents for probing RNA structures in spatially crowded environments, and for assessing the degree of spatial crowding of RNAs in general. Bolstered by the findings, we next applied RISP to measure RNA structural profiles of polyadenylated transcripts in HEK293 cells. We measured the RNA structural profiles with standard probe NAIN3 and the smallest acylating reagent AcIm, which showed relatively higher loop-stem selectivity compared to AcIN3 in rRNAs (Fig. S6). The R-score of each nucleotide for each reagent was calculated via the tuned pipeline (Fig. 4a). The R-score difference of each nucleotide between AcIm and NAIN3 was evaluated by P-values via t-test. Based on the profiles, we assessed the transcriptome-wide signal coverage of each reagent by calculating the number of nucleotides with efficient signal (R-score≥ 0.1) per kilobase. AcIm yielded 80% higher coverage, yielding 210 nt per kb compared to NAIN3 (119 nt per kb) (Fig. 4b). Examination of the 1500 transcripts that contained statistically different sites for the two probes (P<0.05) revealed that 1279 transcripts showed AcIm-dominant signal while only 98 transcripts carried NAIN3 dominant sites (Fig. 4c), indicating better access of the small probe to protein-interacting sites or to tightly folded RNA structure. For the nucleotides showing significant differences between the probes (P<0.05), 44.6% of 2’-OH groups were only probed by AcIm with efficient signal and 54% of them showed large differences, with differential R-score (AcIm vs. NAIN3) ≥ 0.2. In contrast, the 2’-OH groups more efficiently probed by NAIN3 were a much smaller 3.1% of all nucleotides (Fig. 4d). These findings further demonstrated the generality of enhanced intracellular probing ability of a small-sized acylating reagent in transcriptome-wide RNA analyses.

**Figure 4.**
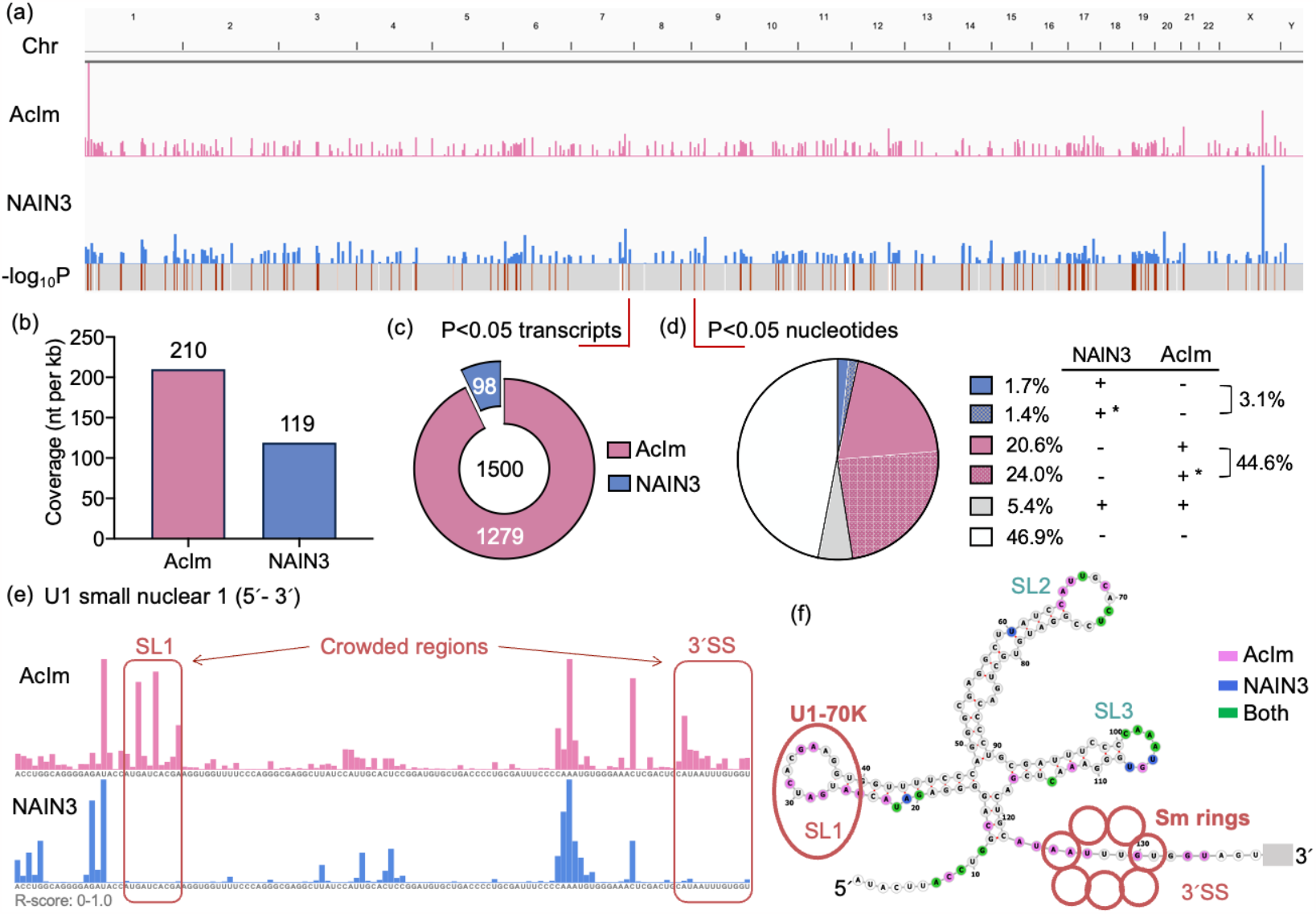
RISP enables enhanced structure probing of RNAs transcriptome-wide. (a) UCSC tracks showing the mean R-scores (y-axis) probed by AcIm (pink) and NAIN3 (blue) in live HEK293 cells mapped along the human chromosomes (*top*) and -Log10*P*-value of Welch *t*-test (*bottom*) of both probes indicating the nucleotides with significant difference (P<0.05). Mean R-scores are scaled from 0 to 1. (b) Structural coverage of AcIm and NAIN3 reagents in polyA+ transcriptome RNAs. The coverage was evaluated by the number of efficiently probed nucleotides per kilobase of sequence. (c) Fractions of transcripts showing AcIm-dominant (1279) and NAIN3-dominant (98) signals in total transcripts (1500) showing significant difference (P<0.05). (d) Distribution of total nucleotides showing significant difference (P<0.05) including only AcIm probed (pink), only NAIN3 probed (blue), both probed (grey) and both unprobed (white). Nucleotide with R-score over 0.1 was efficiently probed by reagents. Nucleotides with differential R-score larger than 0.2 were defined as dominant nucleotides, denoted with “*”. (e) UCSC tracks showing mean R-score of AcIm (pink) and NAIN3 (blue) at an RNA locus in U1 small nuclear 1 RNA (RNU1-1 snRNA), documenting improved structure mapping of small acylating probe AcIm. The regions specifically probed by AcIm are boxed in red, referring to the folding structure as stem-loop 1 (SL1) and 3’ single-strand (3’SS). (f) Mean R-score of AcIm and NAIN3 in (e) mapped onto consensus structural models of U1 snRNA. Only AcIm probed nucleotides are shown in pink; only NAIN3 probed nucleotides in blue; both probed nucleotides are shown in green. SL1 and 3’SS closely interact with U1-70K protein and Sm protein rings, respectively.

We more closely investigated specific transcripts showing signal differences between the small and standard reagents, focusing on cases known to undergo RNA-protein interactions. We examined the U small nuclear RNAs (snRNA) relating to the spliceosome, which contain polyadenylated sites within their annotated 3’ ends in HEK293 cells (Fig. S7). The detected R-scores were mapped to consensus secondary structures. For U1 small nuclear 1 RNA (RNU1-1 snRNA) we found two clear regions (SL1 and 3’SS, see Fig. 4f) showing only AcIm signals. These two regions of the snRNA are known to associate with binding proteins U1-70K and Sm rings (Fig. 4e-f).

We hypothesized that the probing difference of AcIm and NAIN3 was due to a close RNA-protein interaction in SL1 and 3’SS region, which showed a closest distance of 2’-OH to nearby proteins less than 5Å by examining the U1 snRNP crystal structure (Fig. S8a). Although the 5’SS and SL2 regions probed by both AcIm and NAIN3 also interact with U1-C and U1-A proteins, the crystal structure exhibited more spacious interaction between RNA and proteins consistent with the observed reaction of the larger-sized probe (Fig. S8b-c). We also examined other snRNAs (RNU5A-1, RNU4-2, RNU6-7) in conjunction with the crystal structures of their snRNPs in the spliceosome (Fig. S9). The results indicated similar advanced ability of probing nucleotides for AcIm in crowded spaces in comparison to NAIN3.

### RISP enables m^6^A identification in crowded spaces

*N*^6^-Methyladenosine (m^6^A) has been identified as the most abundant and conserved internal posttranscriptional modification within eukaryotic mRNAs (33). The methylation occurs in the consensus sequence DRACH, where the central A is methylated (33). Previous work has identified a specific structure signature for m^6^A modification via NAIN3 based icSHAPE probing in transcriptome RNAs, revealing a stronger icSHAPE reactivity surrounding the m^6^A, possibly due to the destabilization of RNA helices (16). Since the AcIm reagent can probe crowded spaces that are inaccessible to NAIN3, we wondered whether the connection between the probed structure signal and m^6^A modification occurs in crowded regions as well. To test this, we mapped the regions showing significant differences between AcIm and NAIN3 signal (P<0.05) with the published m^6^A database to evaluate the RNA structure profiles of the two reagents around known m^6^A sites (Fig. 5a). We found that 51% of mapped m^6^A sites were efficiently probed by both reagents (suggesting less crowded space), while 44% of the remaining m6A sites were only detected by AcIm (assigned as crowded spaces) (Fig. S10a). By examining the averaging R-score of total probed m^6^A motifs, we observed superior signals of AcIm in both crowded and less-crowded regions in comparison to the standard-sized reagent NAIN3 (Fig. 5b-c, S10b-c). The averaged reactivity pattern revealed a relatively low reactivity at m^6^A and downstream nucleotides, and a strikingly higher reactivity at the position 5’ adjacent to the m^6^A modification (Fig. 5b-c, S10b-c). Notably, previous parallel analysis of RNA structure (PARS) study of isolated human m^6^A-containing RNAs indicated an averaged unpaired structural context for the purine nucleotide immediately upstream of m^6^A and paired for the residues downstream of m^6^A, in accordance with our reactivity observation.

**Figure 5.**
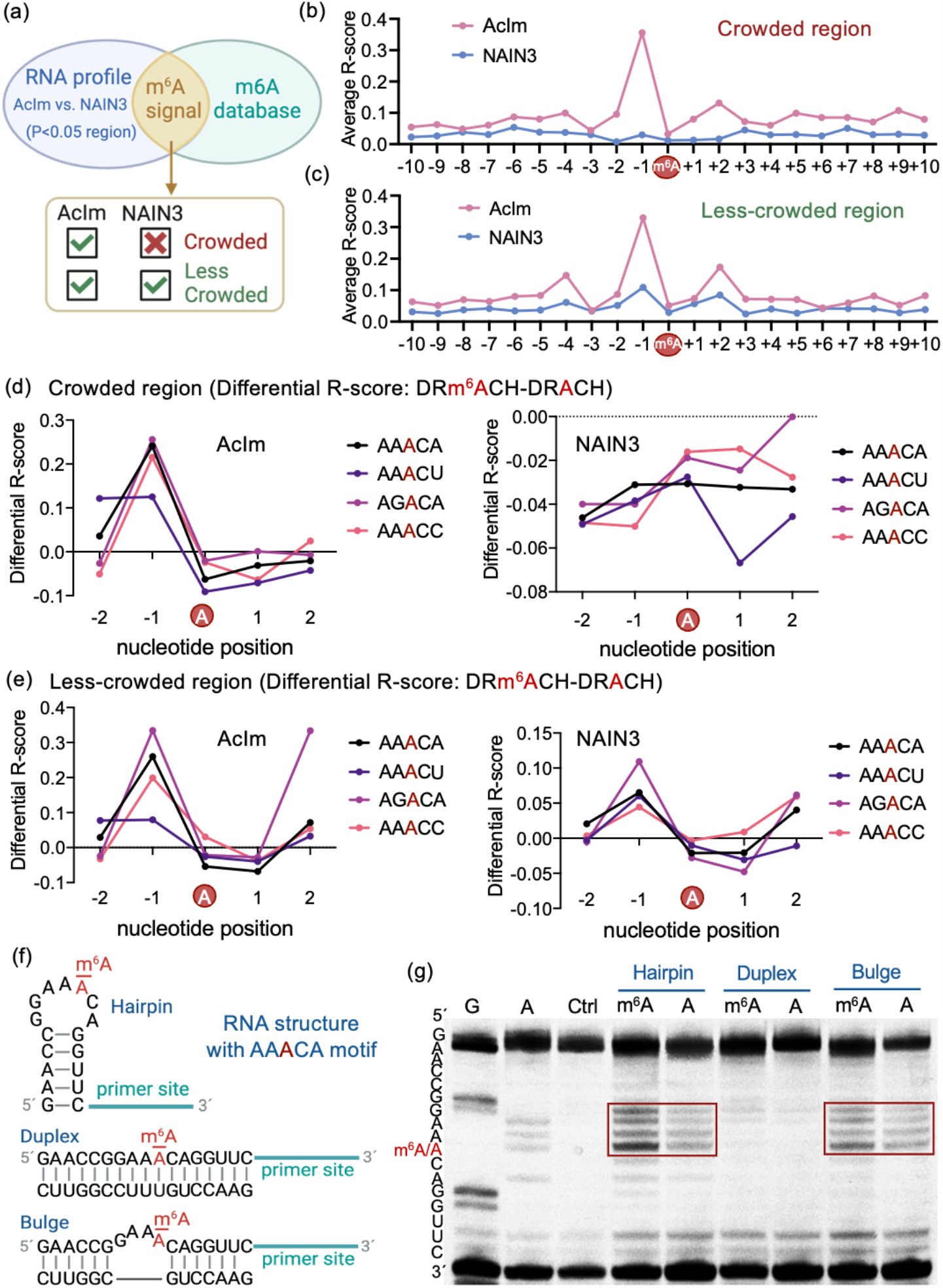
RISP enables N^6^-Methyladenosine (m^6^A) identification in crowded spaces of transcriptome RNAs in living cells. (a) Schematic of mapping transcriptome RNA profiles showing significant difference (P<0.05) for AcIm and NAIN3 within methylation sites defined by the published m^6^A database. Sites that were only probed by AcIm were scored as crowded regions, and regions probed by both reagents but with different signal intensity regarded as less-crowded spaces. (b-c) Average R-score of the region with 10nt upstream and downstream of m^6^A site probed by AcIm and NAIN3 in crowded regions (b) and less-crowded regions (c). (d-e) Differential R-score for m^6^A methylated versus non-methylated sites with the same underlying sequence motifs in crowded regions (d) and less-crowded regions (e). Average R-scores from unmodified motifs were subtracted from m6A-modified motifs. The examined sequences (AAACA, AAACU, AGACA, AAACC) were the top-ranked motifs in our mapping results (Fig. S10d-e). (f-g) In vitro tests of acylation of methylated vs. unmethylated synthetic RNA oligonucleotides of defined sequence/structure using AcIm reagent. (f) Sequences and structures of tested RNAs in the presence and absence of m^6^A modification. RNAs were constructed to form hairpin, duplex and bulge structure containing top ranked m^6^A motif AAACA found in the sequencing data. (g) Gel analysis of reverse transcriptase (RT) stops for designed RNA structures reacting with AcIm, revealing enhanced reactivity at positions 5’ adjacent to m^6^A site, consistent with intracellular data. Note that RT stops (and corresponding bands) occur at the nucleotide immediately 3’ to the reacted position.

Zooming in beyond averaged m^6^A motifs (DRACH), we investigated whether the reactivity pattern differs between individual sequences within the motif by comparing the signals of each detected m^6^A sequence. The results showed no significant difference among the identified m^6^A motifs, suggesting very similar structural contexts that enhance acylation of the nucleotide immediately preceding m^6^A (Figs. S10d-e). To further confirm that the observed signal pattern is specific to m^6^A modification, we compared the R-score of AcIm and NAIN3 at m^6^A-motified vs. unmodified instances of the top four abundant m^6^A motifs in crowded and less-crowded regions (Fig. 5d-e, Fig. S11). The higher differential R-score of 5’ adjacent nucleotide for both reagents in less-crowded regions further supported the specificity of stronger reactivity relating to the m^6^A modification, and again showing the enhanced probing ability of the small probe, which shows ∽3-5-fold more intense signals for methylation (Fig. 5d-e). Interestingly, the smaller probe provides considerably more pronounced signals for methylation in all DRACH contexts (Fig. 5b-e), suggesting that the signal is generated in a crowded structural environment for the nucleotide immediately preceding m^6^A.

Next, we further explored the origins of how methylation of adenine can cause elevated acylation reactivity of the 5’ adjacent nucleotide. In cells, adenine methylation can be recognized by protein “readers” which could cause crowding in nucleotides near m6A. Alternatively, methylation might cause structural changes relative to unmethylated RNA by influencing RNA biophysics: methylation increases base stacking of adenine in unpaired contexts, and conversely, it destabilizes base pairs involving m6A in duplex contexts. The above analyses compared the average reactivity near m^6^A with different surrounding transcriptome contexts, and did so in the presence of possible protein complexes. To isolate the reactivity effects, we designed isolated RNA sequences containing the common AAACA sequence context (which was top-ranked in our data) to fold into defined hairpin, duplex and bulge loop structures in the presence or absence of N^6^-methyl groups as model *in vitro* contexts lacking protein (Fig. 5f). The m^6^A in the bulge loop was situated between unpaired bases and helix, mimicking the preferential m^6^A cellular context revealed by previous studies (34). The constructed RNA structures were probed with 100 mM AcIm for 15min achieving average single-hit reaction levels, analyzed after reverse transcription to identify the acylation of each nucleotide, with (RT) stops appearing immediately 3’ to the acylation adducts. The results showed similar acylation positions but significantly higher reactivity of the adjacent nucleotide 5’ to methyl modification in both hairpin and bulge structures (Fig. 5g), revealing that methylation changes the local loop accessibility/conformation but minimally affects the overall folded structure (35). In contrast, for an m^6^A context in a double-stranded structure we observed little acylation for both modified and unmodified sequences (Fig. 5g), indicating that methylation in stable stems does not alter duplex structure (36). On further examining the solution and crystal structures of RNAs with and without methylated adenine (Fig. S12), we hypothesize that N6 methylation may increase the stacking of adenine with the base immediately 5’ to the modification in loop structures, holding the 2’-OH group of the latter in a more exposed environment on average.

## Discussion

RNAs can fold into intricate and compact 3D structures and are often accompanied by regulatory interactions with other macromolecules in cells. Many functional non-coding RNAs such as ribosomal RNAs and spliceosomal snRNAs function in highly assembled ribonucleoprotein complexes (2). Further, approximately 95% of cellular mRNAs have identified binding proteins (19). The most frequently used chemical probing reagents map RNA structure by accessing and reacting with the 2’-OH group in RNAs. Although efficient, structural coverage obtained with current probes is significantly limited by the inaccessibility of the reagents to occluded structure and environment. This study establishes RISP for mapping RNA structure with close spatial contacts, taking advantage of differentially sized acylating reagents (AcIm/AcIN3/NAIN3) and a comparative workflow to access the regions where RNAs are closely interacting with proteins and with nearby RNA structure. Compared with a considerably larger standard probe, our data demonstrate substantially greater structural probing ability for the small AcIm probe within tunnels and crevices between and underlying proteins in the human 80S ribosome. Thus, RNA infrastructure that was previously functionally invisible can be probed with reagents such as AcIm that can physically reach into spaces too small for standard probes. As a result, AcIm profiling can provide considerably expanded coverage of cellular RNAs due to this improved access. While a different class of RNA structural probe, dimethylsulfate (DMS), is a similarly small molecule as AcIm, there is currently no larger alkylating agent that has been developed for comparison such as in RISP. In addition, DMS provides information only for cytosine and adenine bases, whereas acylation probes can target essentially every accessible nucleotide in the transcriptome.

In addition to providing greater depth of RNA folding information, the RISP approach can help to infer distances and tunnel spaces of nucleotides close to nearby macromolecules. In our model ribosome analysis, AcIm/AcIN3 show significant probing frequency for nucleotides with average distance less than 5Å to the closest proteins, whereas NAIN3 gives minimal signal in the same ranges (Fig. 2b). Similarly, in our ribosomal tunnel space analysis, acetyl probes accessed nucleotides with tunnel radius down to 1Å which prevents NAIN3 access (Fig. 3c, S5b). Thus, comparing the profiles of AcIm/AcIN3 and NAIN3, the user can infer that regions that predominated by the small-sized probes are likely to have macromolecule contacts within 5Å distance. Such information could help to improve the 3D structure modeling of RNA complexes in the setting of living cells.

We also applied RISP transcriptome-wide to polyadenylated RNAs in cells. We find that AcIm yields 80% greater structural coverage than NAIN3 (Fig. 4b) among the ∽2000 transcripts containing sufficient data for comparative analysis. Our data show that ∽85% of these transcripts exhibited superior probing coverage with the smaller AcIm (Fig. 4c). The new results confirm that small acetyl reagents provide structural data with broader coverage and higher signal, adding considerable amounts of structural information to existing in-cell acylation maps of the human transcriptome. In addition, the mRNA regions with greatly different signals between the small and standard-sized probes may be considered candidates for known and unknown protein interactions.

In addition to providing data on folding and potential protein interactions, our RISP analysis also shows that the small reagent AcIm can provide clear signals and structural details for m^6^A methylation sites of RNAs in their native cellular setting. We find that AcIm provides higher signal over background than reagent NAIN3 in probing methylation generally, and yields an even greater advantage in crowded spaces where no efficient methylation signal is produced by the latter. The new data support previous studies that m^6^A methylation primarily occurs at sites that contain unpaired motifs (16, 37). The observed m^6^A signal pattern shows strikingly stronger reactivity at the adjacent position 5’ to the m^6^A in the DRACH motif. Interestingly, previous nuclease structural mapping of immunoprecipitated human m^6^A RNAs revealed a structural transition from unpaired to paired surrounding m^6^A over the averaged motif, showing that the purine nucleotide immediately upstream of m^6^A has a high probability of being unpaired (34). This may explain our observation of the high R-score for the 5’ adjacent nucleotide to the modification in our transcriptome profile. Our studies of isolated synthetic RNAs with and without an N^6^-methyl modification confirmed similar acylation behavior, with elevated reactivity 5’ to m^6^A in vitro (Fig. 6). This confirms that the acylation signal we observed in HEK293 cells for m6A is not a result of protein interactions. We hypothesize that the methylation of adenine might affect local loop confirmation by increasing the stacking with the 5’ adjacent purine base to increase the steric exposure of the 2’-OH group for enhanced reagent accessibility (Fig. S12). Several approaches have been developed for assessing m^6^A sites in cells. Immunoprecipitation methods (m^6^A-seq, miCLIP, and m^6^A-LAIC-seq) and enzyme-assisted approaches (DART-seq, MAZTER-seq, m^6^A-REF-seq, m^6^A-label-seq, m^6^A-SAC-seq) have been reported (38, 39). Although these methods are efficient for detection and provide quantitative analysis of m^6^A, the use of the simple AcIm probe method requires no engineered protein constructs for m^6^A site detection directly *in cellulo*. Given the ready commercial availability of the probe, our RISP method could be a useful addition to current methods for analysis of m^6^A sites in the transcriptome.

## Data availability

Our RNA structure mapping data will be posted on the GEO database for future reference. Datasets from previous publications have reported on the details of the experiments and the corresponding analyses.

## Acknowledgement

We thank the U.S. National Institutes of Health (GM145357) for support.

