## Supplemental Information for "RNA Infrastructure Profiling Illuminates Transcriptome Structure in Crowded Spaces"

**Table of contents**

- Supplementary Table 1. List of oligonucleotides used in this work S2
- Supplementary Table 2. List of reagents and materials S2-S3
- Supplementary Figures S4-S17
- Experimental Procedures S18-S20
- References S20-21

**Table S1. List of oligonucleotides used in this work**

| Name | Sequence (left to right: 5' to 3') |
|---|---|
| $m^6$A-RNA-chimeric | rGrArArCrCrGrGrArA/iN6Me-rA/rCrArGrGrUrUrC GACAGAAATAAAAAAGGGA |
| A-RNA-chimeric | rGrArArCrCrGrGrArArArCrArGrGrUrUrC GACAGAAATAAAAAAGGGA |
| Helper RNA-duplex | rGrArArCrCrUrGrUrUrUrCrCrGrGrUrUrC |
| Helper RNA-bulge | rGrArArCrCrUrGrCrGrGrUrUrC |
| A/$m^6$A-RNA RT primer | /FAM/ TCCCTTTTTTATTTCTG |
| Human 5S rRNA RT primer | /Cy5/ AAAGCCTACAGCACCCGGTAT |

**Table 2. List of all reagents and materials**

| REAGENTS | SOURCE | IDENTIFIER |
|---|---|---|
| **Reagents and Enzymes** | | |
| 5M NaCl | Invitrogen™ | # AM9760G |
| 1M $MgCl_2$ | Invitrogen™ | # AM9530G |
| 0.5M MOPS buffer, pH7.5 | Thermo Scientific™ | # J62839-AE |
| UltraPure DTT | Invitrogen™ | # 15508013 |
| NTP Set (100 mM Solution) | Thermo Scientific™ | # R0481 |
| dNTP mix (10 mM each) | Thermo Scientific™ | # R0194 |
| SYBR Green I nucleic acid gel stain (10,000×) | Invitrogen™ | # S7567 |
| SYBR Gold Nucleic Acid gel stain (10,000X) | Thermo Scientific | # S11494 |
| SequaGel - UreaGel Concentrate | National diagnostics | # EC-830 |
| SequaGel - UreaGel Diluent | National diagnostics | # EC-840 |
| SequaGel - UreaGel Buffer | National diagnostics | # EC-835 |
| Agarose | Fisher Scientific | # BP1356500 |
| Ammonium Persulfate | Thermo Scientific™ | # 17874 |
| N, N, N′, N′-Tetramethyl ethylenediamine | Sigma-Aldrich | # 110732 |
| Topvision agarose | Thermo Scientific™ | # R0491 |
| 10xTBE buffer | KD Medical | # RGF-3330 |
| RNaseOUT™ Recombinant Ribonuclease Inhibitor | Invitrogen™ | # 10777019 |
| RiboLock RNase inhibitor | Thermo Scientific™ | # EO0382 |
| FastAP thermosensitive alkaline phosphatase | Thermo Scientific™ | # EF0652 |
| RQ1 RNase-Free DNase I | Promega | # M6101 |
| T4 polynucleotide kinase | New England BioLabs | # M0201S |
| T4 RNA ligase 1 | New England BioLabs | # M0204S |
| RNase cocktail enzyme mix | Invitrogen™ | # AM2286 |
| RNase H | Thermo Scientific™ | # EN0201 |
| CircLigase II ssDNA ligase | Epicentre | # CL9025K |
| SuperScript™ III Reverse Transcriptase | Invitrogen™ | # 18080044 |
| Trizol LS Reagent | Thermo Scientific | # 10296028 |
| Dulbecco’s Modified Eagle Medium (DMEM) | gibco | # 11995-065 |
| 1xPBS, pH=7.4 | gibco | # 10010-023 |
| UltraPure DNase/RNase-free Distilled water | Thermo Scientific | # 10977023 |

| | | |
|---|---|---|
| Glycogen, RNA grade | Thermo Scientific | # R0551 |
| 3 M sodium acetate, pH=5.5 | Thermo Scientific | # AM9740 |
| 96% Ethanol | Fisher | # BP8202-500 |
| Fetal bovine serum (FBS) | gibco | # 26140-079 |
| RNA Gel loading dye (2x) | Thermo Scientific | # R0641 |
| RNA Fragmentation Reagents | Thermo Scientific | # AM8740 |
| **Commercial kits** | | |
| Corning Costar Spin-X centrifuge filters, 0.45 µm | Millipore-Sigma | # CLS8162 |
| RNA clean-up and concentrator-5 column | Zymo Research | # R1016 |
| DNA clean-up and concentrator-5 column | Zymo Research | # D4014 |
| Amicon Ultra-0.5 10K-Centrifugal Filter Unit | Millipore-Sigma | # UFC500396 |
| Phusion high-fidelity (HF) PCR master mix | New England BioLabs | # M0531S |
| Q5 hot start high fidelity PCR master mix | New England BioLabs | # M0494S |
| Poly(A)Purist MAG kit | Thermo Scientific | # AM1922 |
| MiniElute gel extraction Kit | QIAGEN | # 28604 |
| Quick-RNA Midiprep kit | Zymo Research | # R1056 |
| **Chemicals** | | |
| Azidoacetic Acid | TCI | # A3079 |
| 1,1'-carbonyldiimidazole (CDI) | Sigma-Aldrich | #115533 |
| DMSO | ACROS Organics | # B0532976 |
| Acetic acid | Glacial Fisher Scientific | # A38-212 |
| Chloroform | Fisher Scientific | #AC610281000 |
| **Cell lines** | | |
| HEK293 | ATCC | #CRL-1573 |
| **Software and online resource** | | |
| RBRP bioinformatics pipeline | Github | https://github.com/linglanfang/RBRP |
| icSHAPE bioinformatics pipeline | Github | https://github.com/qczhang/icSHAPE |
| UCSC Genome Browser | UCSC Genomics Ins. | https://genome.ucsc.edu/ |
| Integrative Genomics Viewer (IGV) | Broad Institute | https://software.broadinstitute.org |
| wiggletools | Github | https://github.com/Ensembl/WiggleTools |
| Bedtools | Github | https://github.com/arq5x/bedtools2 |
| SeqKit | Github | https://github.com/shenwei356/seqkit |
| MOLEonline | ELIXIR-CZ research | https://mole.upol.cz/ |
| DirectRMDB | Zhang et. al NAR, 2022 | http://www.rnamd.org/directRMDB/ |
| ImageJ | NIH | N/A |
| PyMOL | Schrödinger | N/A |
| GraphPad Prism 9 | GraphPad Prism | N/A |
| MestReNova | Mestrelab | N/A |
| BioRender | BioRender.com | N/A |

### Supplementary Figures

(a) NAIN3

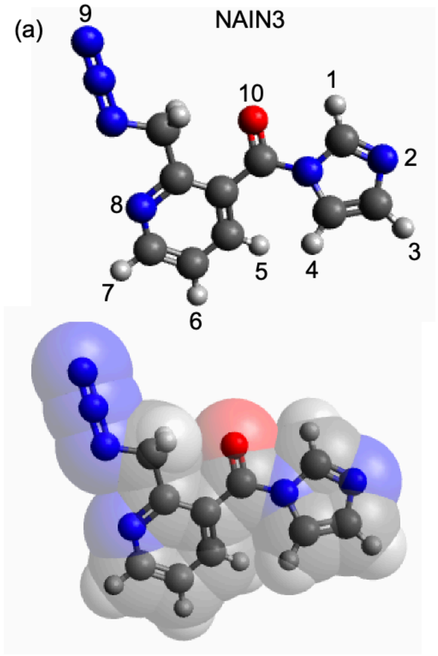


(b) NAIN3

| Atom-to-Atom | Distance (Å) |
|---|---|
| 1-3 | 4.167 |
| 1-4 | 4.219 |
| 1-5 | 4.910 |
| 1-6 | 7.201 |
| 1-7 | 8.101 |
| 1-8 | 6.506 |
| 1-9 | 7.438 |
| 2-4 | 3.302 |
| 2-6 | 7.003 |
| 2-7 | 8.465 |
| 2-8 | 7.263 |
| 2-9 | 9.226 |
| 3-9 | 10.62 |

(c) AcIm

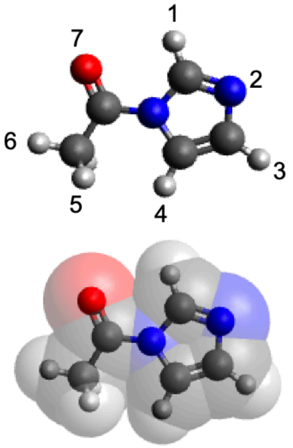


(d) AcIm

| Atom-to-Atom | Distance (Å) | Atom-to-Atom | Distance (Å) |
|---|---|---|---|
| 1-3 | 4.175 | 2-7 | 4.082 |
| 1-4 | 4.225 | 3-5 | 4.926 |
| 1-5 | 4.685 | 3-6 | 6.220 |
| 1-6 | 4.842 | 3-7 | 5.508 |
| 1-7 | 2.591 | 4-6 | 3.780 |
| 2-4 | 3.312 | 4-7 | 4.046 |
| 2-5 | 4.787 | 5-7 | 3.169 |
| 2-6 | 5.601 | 6-7 | 2.559 |

**Figure S1.** Size comparison of small (AcIm) and standard (NAIN3) probes. (a) and (c) Energy-minimized models of NAIN3 and AcIm. Bottom shows Van der Waals surfaces. Black: carbon; blue: nitrogen; red: oxygen; light grey: hydrogen. (b) and (d) Atom-to-atom distances of NAIN3 and AcIm. The displayed largest distance is colored in red, and the shortest distance is in blue.

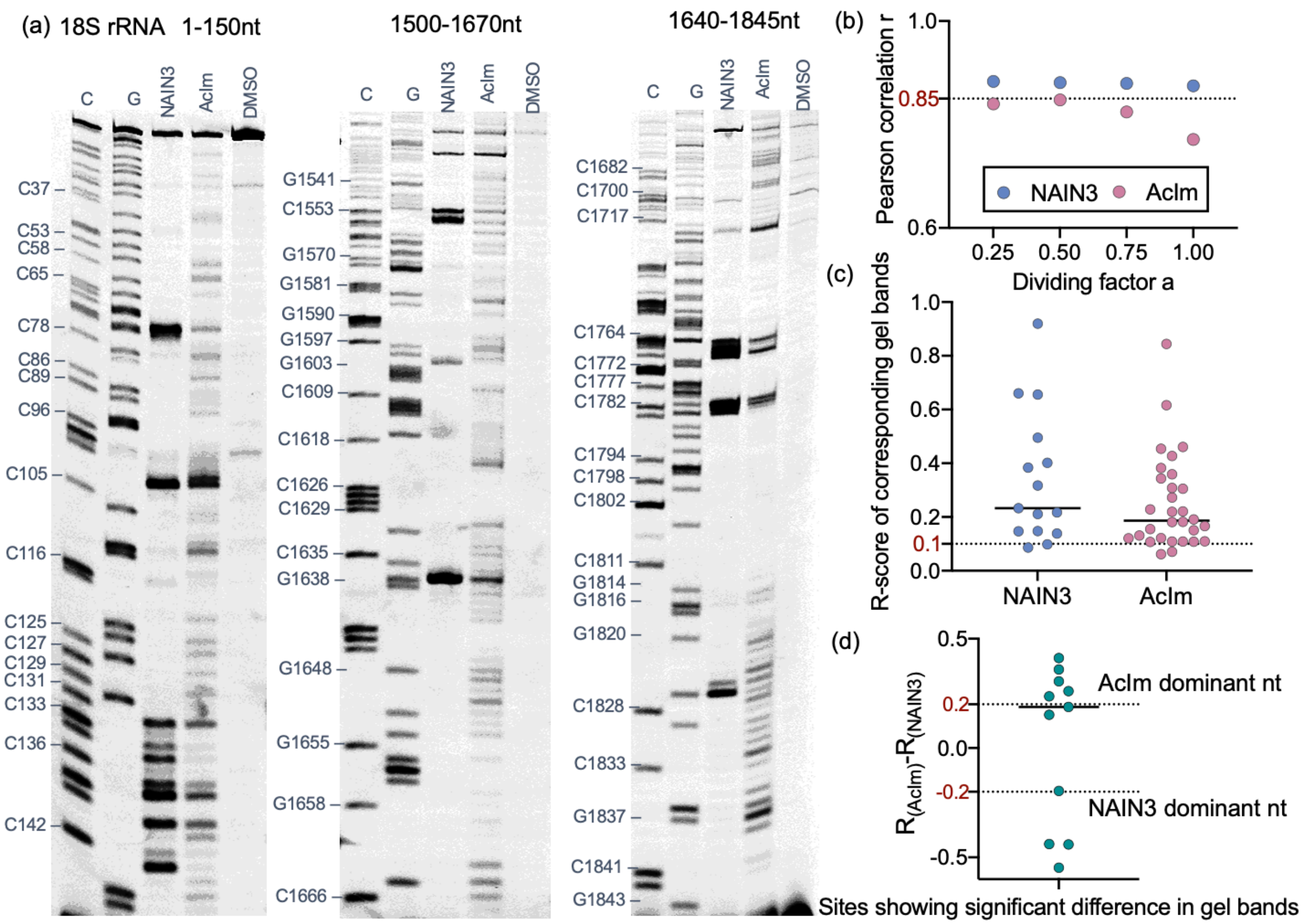


**Figure S2.** Meta-analysis of structural probing gels with calculated R-score as RISP signal for bioinformatics tuning and criteria setting. (a) Structural probing PAGE gels analyzing segments of 18S rRNA in HEK293 cells. The gel data were obtained from our previous work (1). (b) Meta-analysis to adjust the parameters of the bioinformatics pipeline for strongest correlation between manual mapping gels and calculated R-scores. (c-d) Meta-analysis of the RT stops in gels with our sequencing data, setting the threshold of larger than 0.1 as an efficient signal (R-score≥0.1) (c) and the criteria of probe-dominant nucleotides by comparing the differential R-score of small and standard reagent profiles (d). Probe-dominant nucleotides have an absolute differential R-score greater than 0.2 (R-score≥0.2).

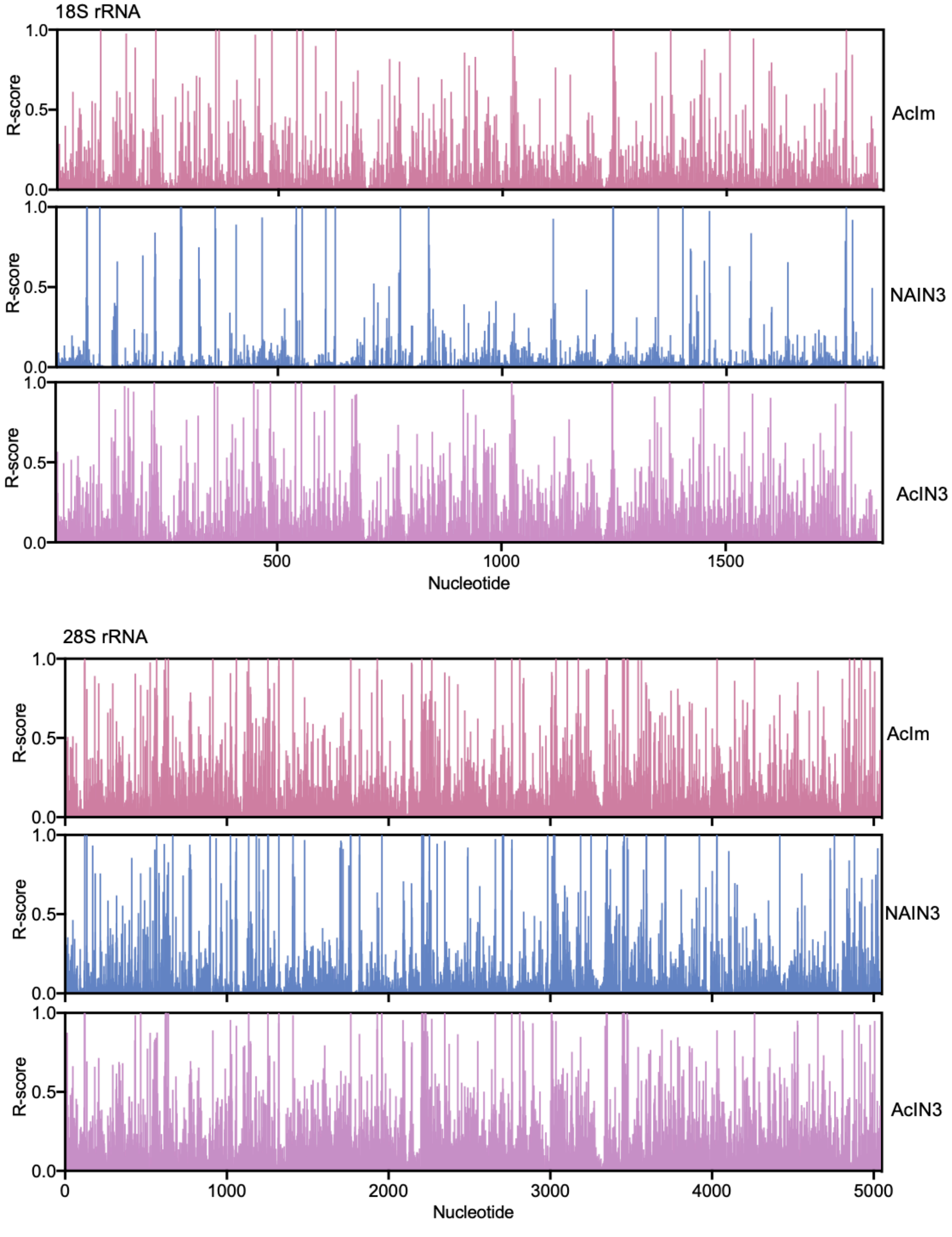


**Figure S3**. Calculated mean R-score of each nucleotide in 18S and 28S rRNA probed by AcIm, AcIN3, and NAIN3 reagents.

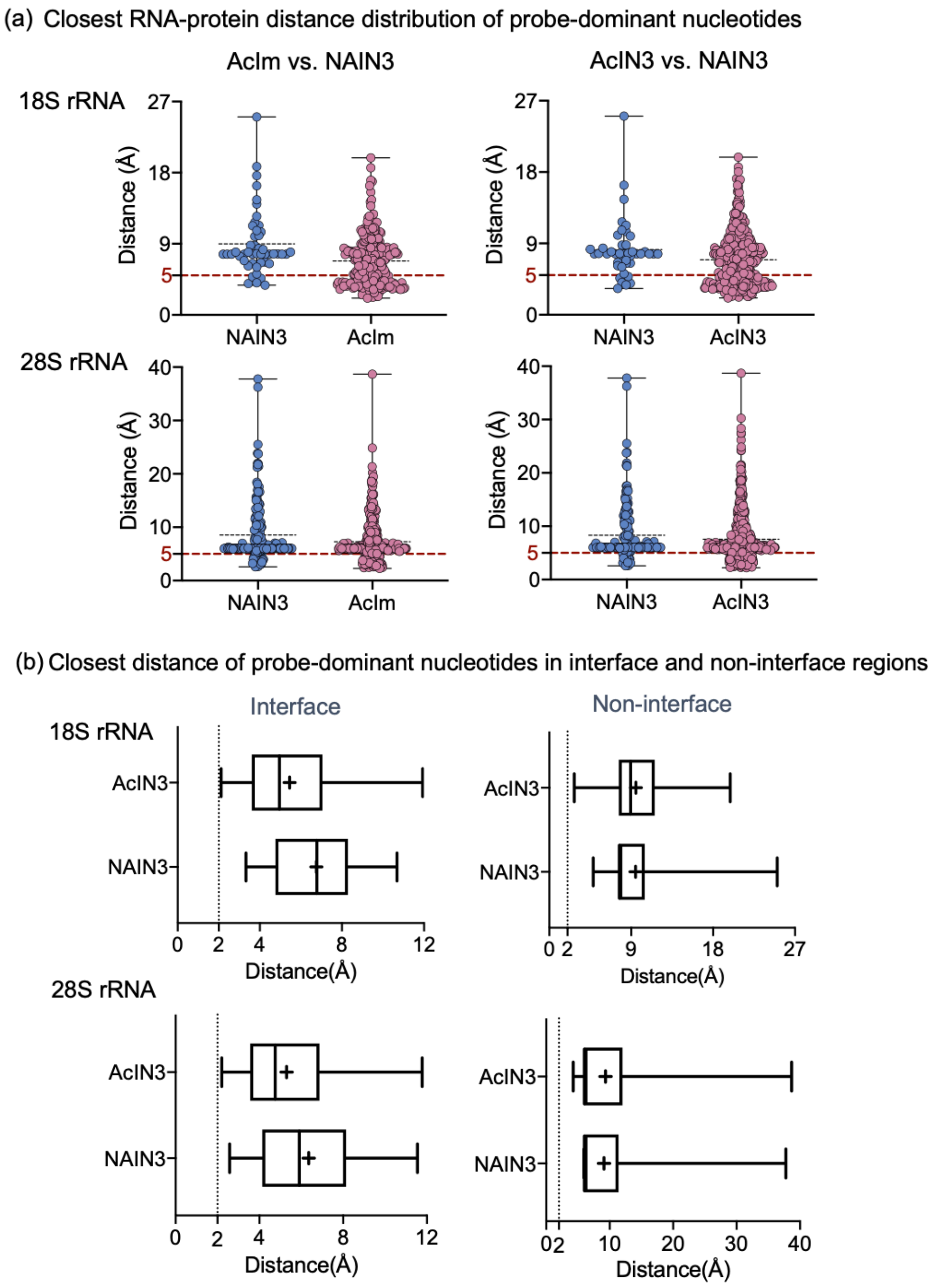


**Figure S4.** (a) Violin plot of the closest RNA-protein distances of dominant nucleotides in 18S and 28S rRNAs for AcIm, AcIN3, and NAIN3, revealing enhanced ability of small reagents (AcIm, AcIN3) in probing nucleotides close to interacting proteins. (b) The closest RNA-protein distance comparison of dominant nucleotide probed by AcIN3 and NAIN3 reagents located in the 18S and 28S rRNA-protein interface and non-interface regions.

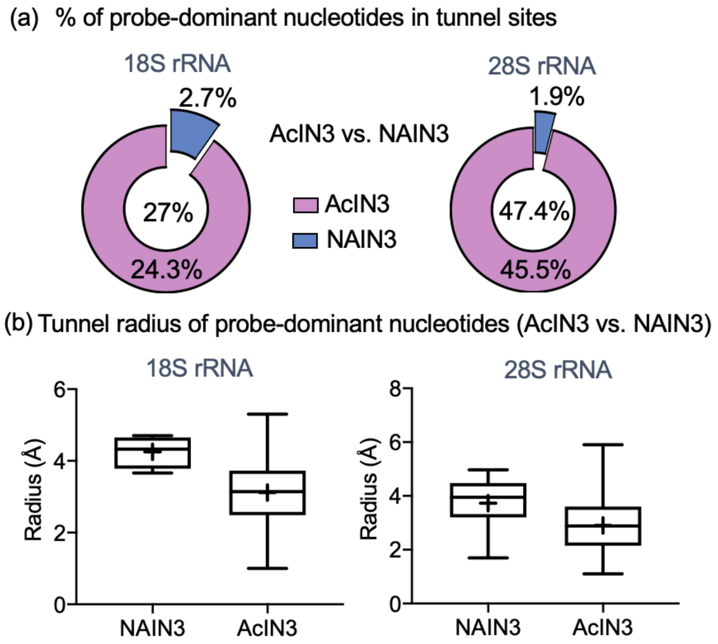


**Figure S5.** Dominance of a small probe in sterically restricted rRNA sites. (a) Percentage of AcIN3- and NAIN3-dominant nucleotides in probed ribosomal RNA tunnel sites for 18S and 28S rRNA. (b) Tunnel radius analysis of AcIN3 and NAIN3 dominant nucleotides in probed RNA tunnel sites for 18S and 28S rRNA.

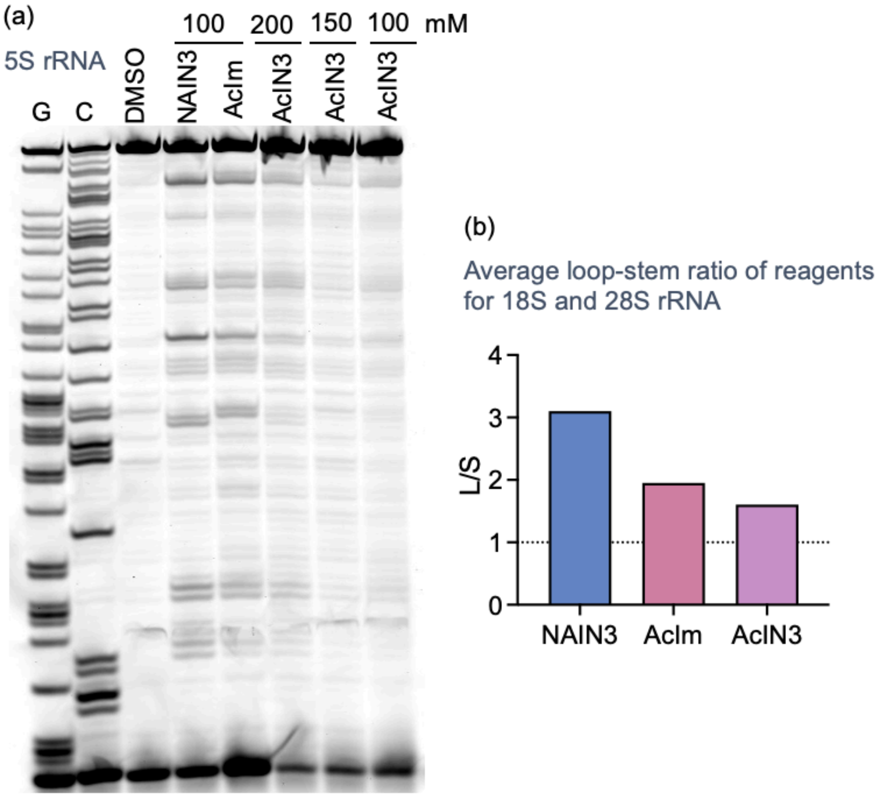


**Figure S6.** (a) Gel analysis of RT stops for 5S rRNA probed by NAIN3, AcIm and AcIN3 at different concentrations. (b) Loop-stem ratio (L/S) evaluated by average R-score of nucleotides in 18S and 28S rRNA. NAIN3 and AcIm show relatively higher loop-stem dynamic range than AcIN3.

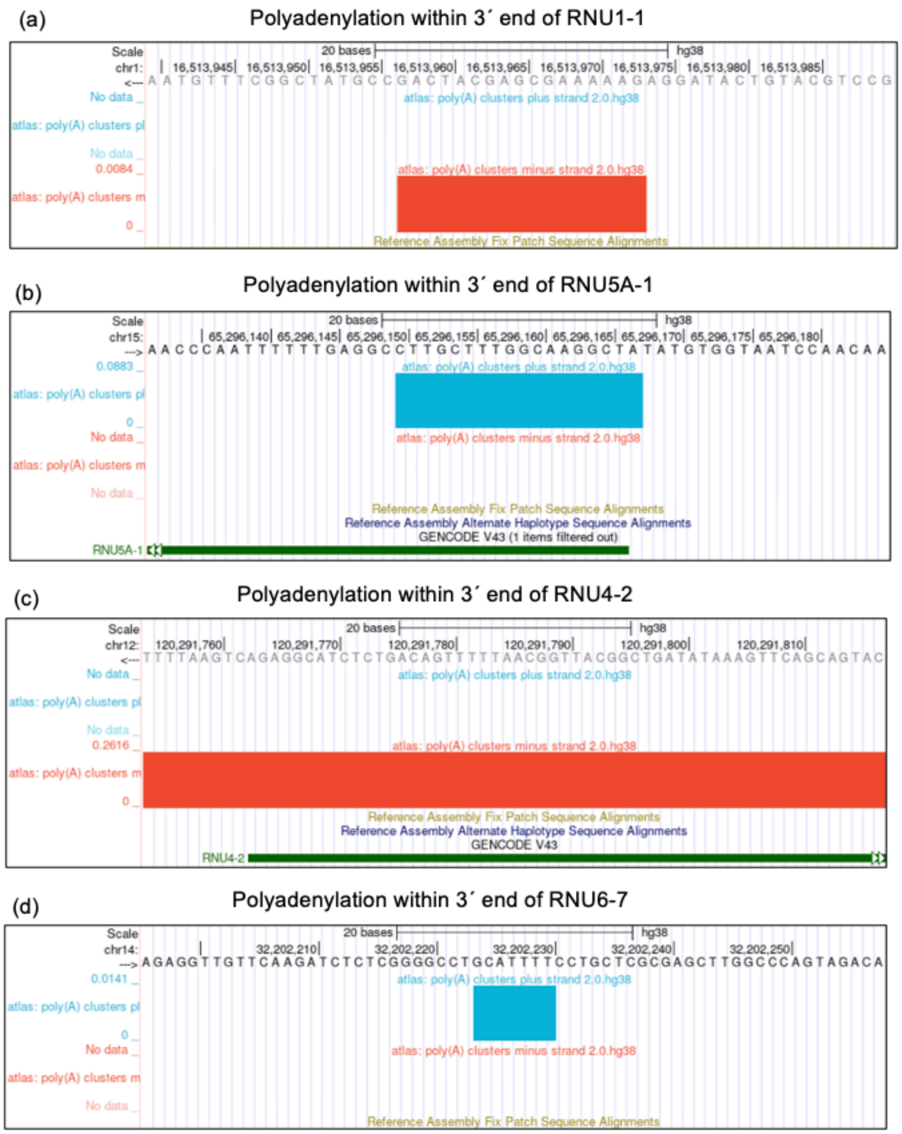


**Figure S7.** Polyadenylation (poly(A)) analysis of U family snRNA associated with spliceosome from published databases. The data show that U snRNAs contain poly(A) site at 3´ ends during their life cycles of processing. (a-d) UCSC track showing Poly(A) sites within the annotated 3´ end of RNU1-1(Chr1: 16513957-16513973; negative strand), RNU5A-1(chr15:65296150-65296167; positive strand), RNU4-2 (chr12:120291754-120291817; negative strand), and RNU6-7 (chr14:32202224-32202230; positive strand). The UCSC tracks were generated with PolyASite 2.01, which was constructed from publicly available human 3´ end sequencing datasets.

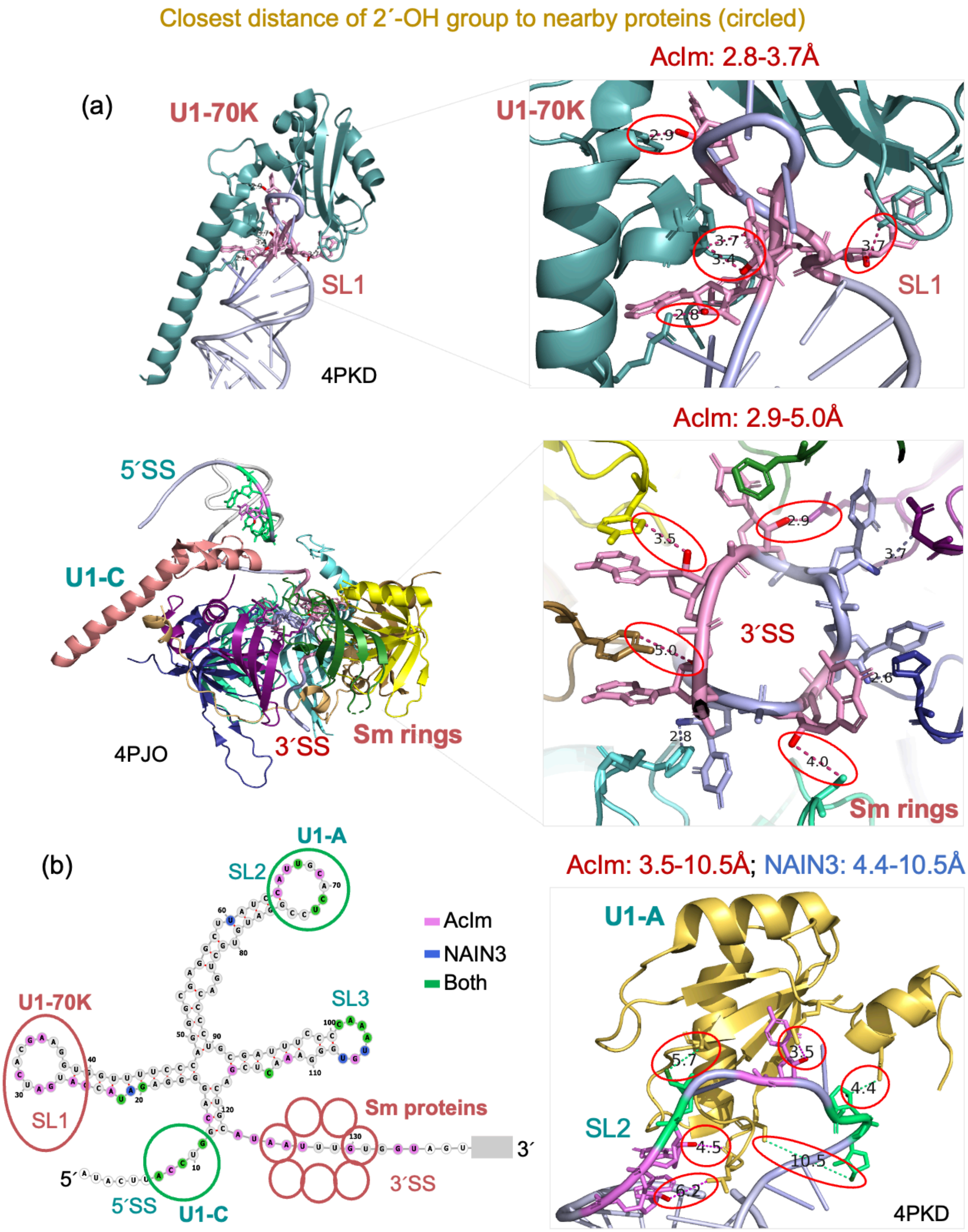
Closest distance of 2´-OH group to nearby proteins (circled)
(a)
U1-70K
SL1
4PKD
AcIm: 2.8-3.7Å
U1-70K
2.9
3.7
3.4
3.7
2.8
SL1
5´SS
U1-C
4PJO
3´SS
Sm rings
AcIm: 2.9-5.0Å
3.5
2.9
3.7
5.0
3´SS
2.6
2.8
4.0
Sm rings
(b)
U1-A
SL2
SL3
U1-70K
SL1
Sm proteins
3´SS
5´
3´
5´SS
U1-C
AcIm
NAIN3
Both
AcIm: 3.5-10.5Å; NAIN3: 4.4-10.5Å
U1-A
5.7
3.5
4.4
SL2
4.5
10.5
6.2
4PKD

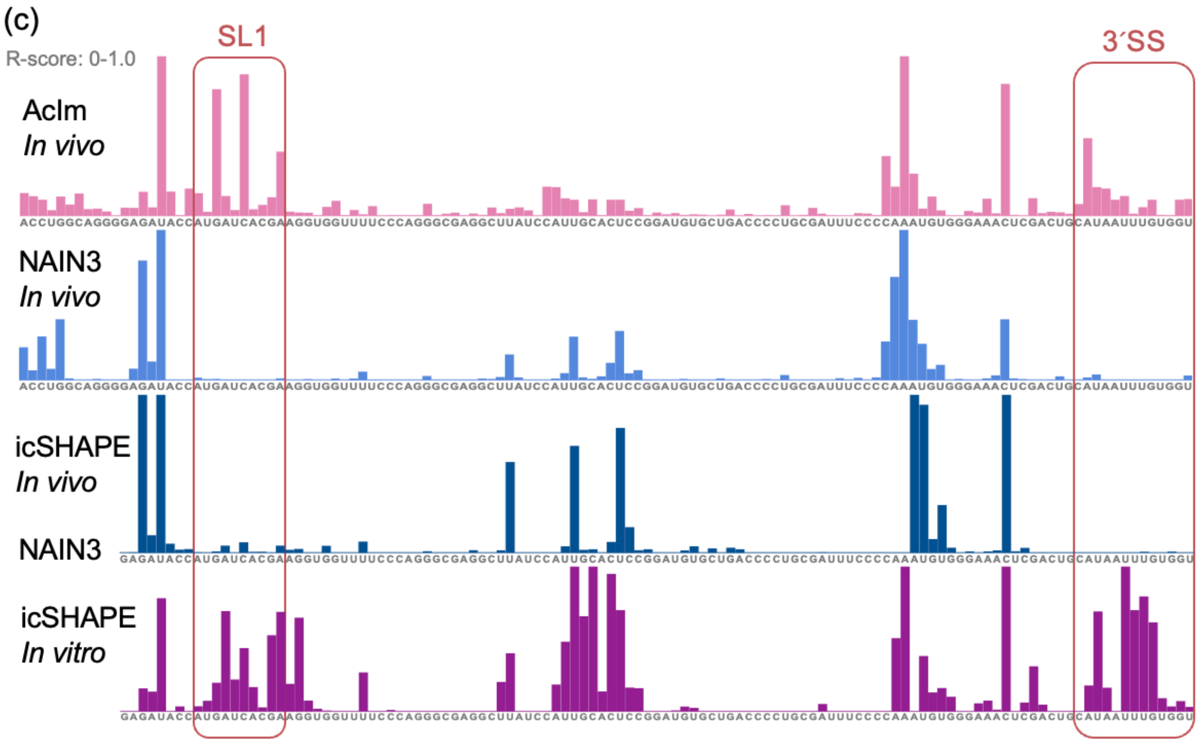


**Figure S8**. Structure-based analysis of close protein-RNA interactions in crowded regions of U1 snRNA. (a) Crystal structure of U1 small nuclear RNA 1 (U1 snRNA) interacting with proteins of U1-70K, Sm rings, U1-A and U1-C proteins. The closest distance of AcIm-reacted 2´-OH to nearby protein heavy atoms (C,N,O) ranges from 2.8 Å to 5 Å in uniquely AcIm-probed regions (SL1 and 3´SS). In both probed region SL2 interacting with U1-A protein, the closest RNA-protein distance for AcIm is ranging from 3.5-10.5Å, whereas for NAIN3 is 4.4-10.5 Å. (b) Secondary structure of U1 snRNA mapping with probed nucleotides and denoting with all interacting proteins (U1-70K, Sm rings, U1-A, U1-C) for each structural region (SL1, 3´SS, SL2, 5´SS). The nucleotides colored in pink were only efficiently probed by AcIm, while those colored in blue were only efficiently probed by NAIN3 and in green were detected by both reagents. PDB: 4PKD, 4PJO. (c) Meta-analysis of reported *in vivo* and *in vitro* icSHAPE data of polyadenylated transcripts produced by NAIN3 reagent (bottom two) with our *in vivo* profiling data yielded by AcIm and NAIN3 (top two). Our profile generated by NAIN3 is consistent with the published *in vivo* structural data, while the RNA profiles probed by AcIm showed similarity with *in vitro* icSHAPE data in the absence of cellular proteins, both detecting the loop (SL1) and single-stranded (3´SS) structures that are tightly associated with U1-70K and Sm protein.

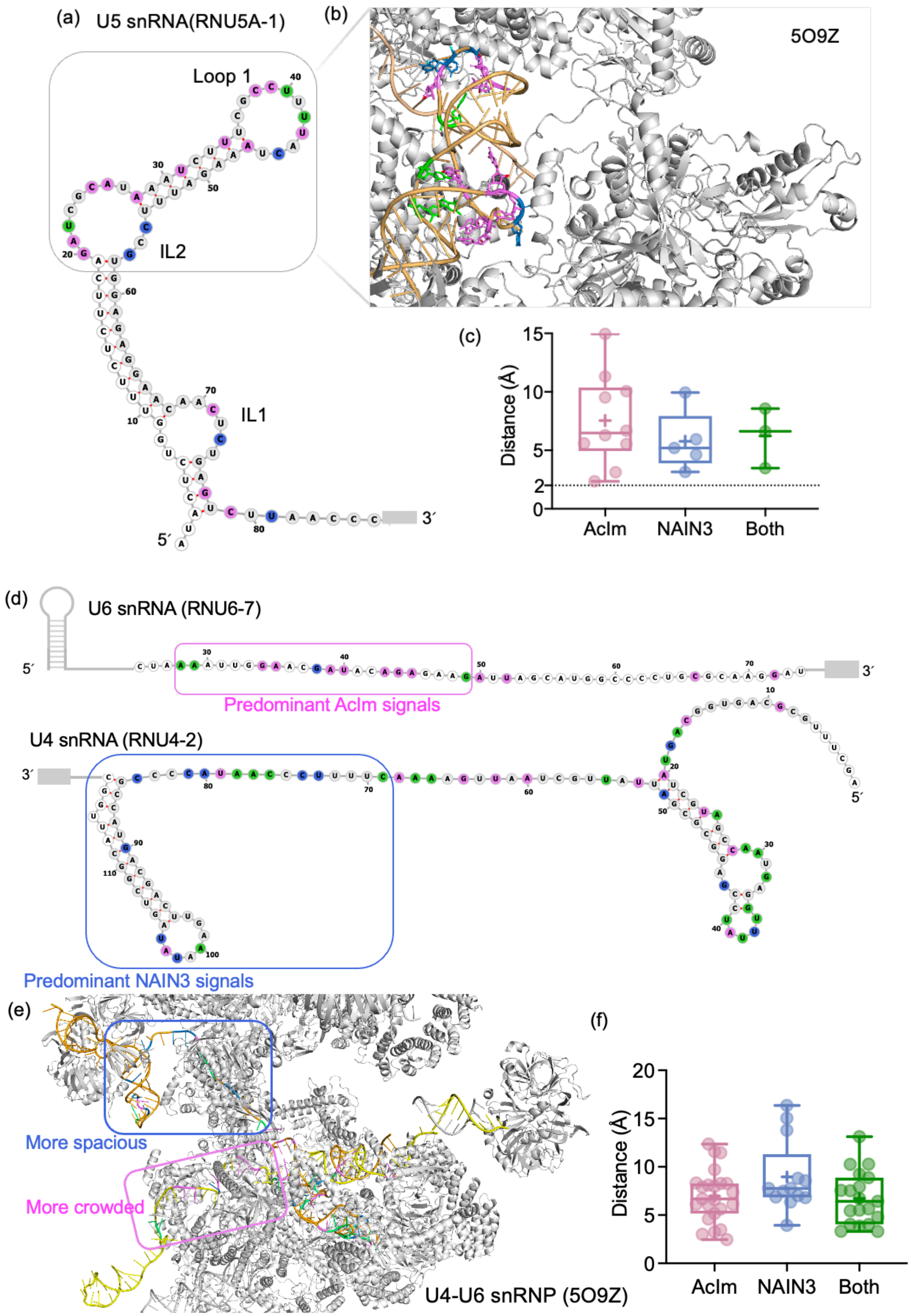
(a) U5 snRNA(RNU5A-1)
Loop 1
IL2
IL1
5´
3´
(b)
5O9Z
(c)
Distance (Å)
AcIm
NAIN3
Both
(d)
U6 snRNA (RNU6-7)
Predominant AcIm signals
U4 snRNA (RNU4-2)
Predominant NAIN3 signals
(e)
More spacious
More crowded
U4-U6 snRNP (5O9Z)
(f)
Distance (Å)
AcIm
NAIN3
Both

**Figure S9.** Structure-based analysis of close protein-RNA interactions of other U snRNAs in spliceosomes. (a) Secondary structure of U5 snRNA (RNU5A-1) mapping with probed nucleotides. Pink: only AcIm probed nucleotides; blue: only NAIN3 probed nucleotides; green: both reagents probed nucleotides. (b) Crystal structure of U5 snRNP with probed nucleotides corresponding to (a). (c) The closest RNA-protein distance of total probed nucleotides corresponding to (a) and (b). (d) Secondary structure of U6 snRNA (RNU6-7) and U4 snRNA (RNU4-2) mapping with probed nucleotides. Pink: only AcIm probed nucleotides; blue: only NAIN3 probed nucleotides; green: both reagents probed nucleotides. (e) Crystal structure of U4 and U6 snRNP with probed nucleotides corresponding to (d). (f) The closest RNA-protein distance of total probed nucleotides corresponding to (d) and (e). PDB: 5O9Z. Note that the profiles of AcIm and NAIN3 represent an averaged structure as a snapshot of snRNPs of spliceosome; it is not expected to be identical to the crystal structure, as the probing covers the RNP in varied states of assembly in the cell. Overall, the results indicate that AcIm can probe more nucleotides in crowded spaces while NAIN3 tends to show dominant signals in more spacious regions.

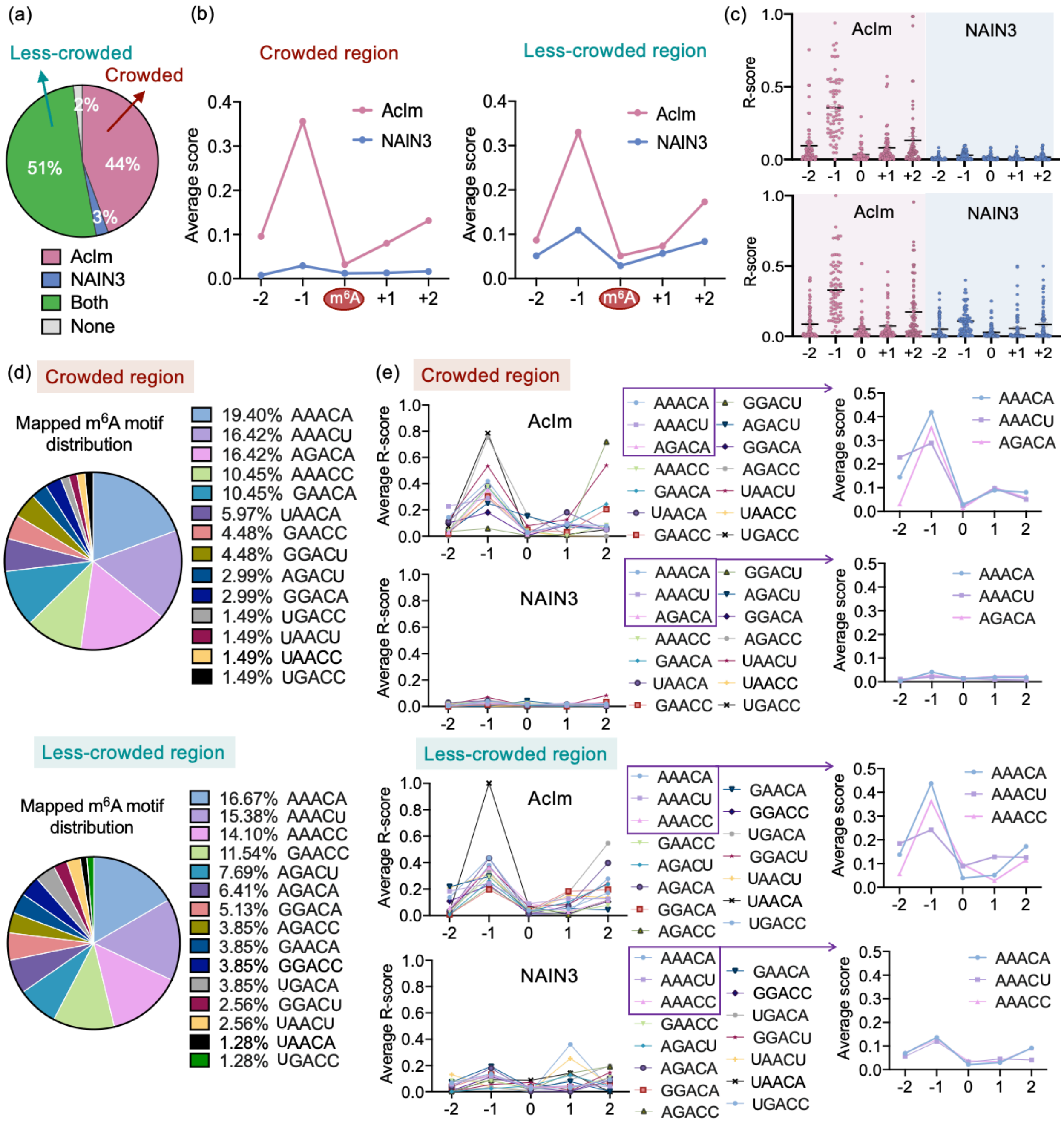


**Figure S10.** AcIm provides improved signals for m6A methylation sites in cellular transcripts. (a) Percentage of total mapped m6A in database probed by AcIm only (pink), NAIN3 only (blue), both agents (green), and unprobed (grey). (b) Average R-score of 5nt m$^6$A motifs probed by AcIm and NAIN3 in crowded regions and less-crowded regions. (c) Scatter plot of R-score of 5nt m6A motifs probed by AcIm (pink) and NAIN3 (blue) in crowd (top) and less-crowd (bottom) regions, corresponding to fig. 5b-c. (d) Distribution of mapped m6A motif sequence in crowd and less crowd regions. (e) Average R-score of each 5nt m6A motif (DRACH) probed by AcIm and NAIN3 in crowded and less-crowded regions, showing similar reactivity pattern for each sequence.

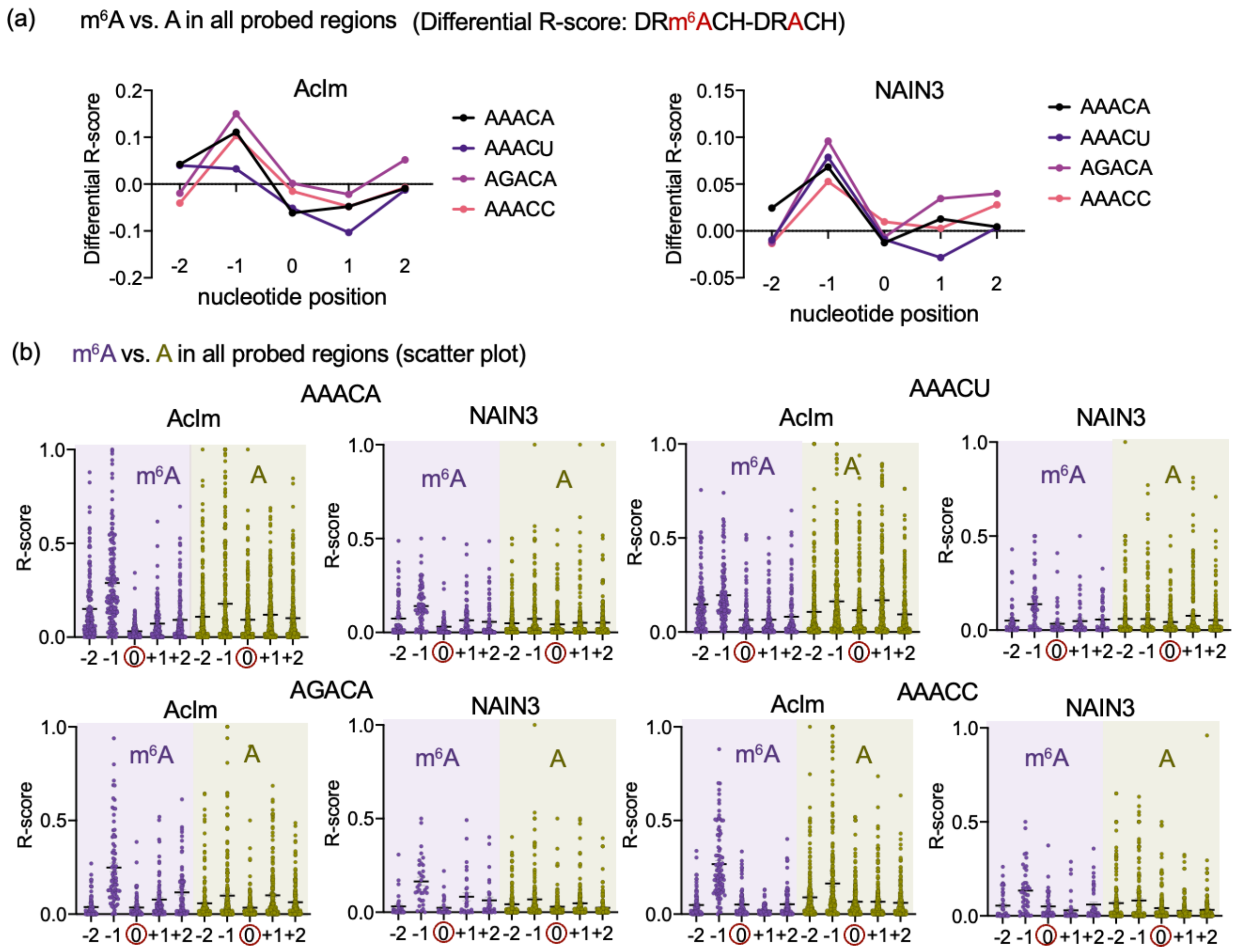


**Figure S11.** Comparison of R-scores of AcIm and NAIN3 at m$^6$A-motified vs. unmodified instances in all probed regions, revealing the observed signal pattern specific to m$^6$A modification. (a) Differential R-score for m$^6$A methylated versus non-methylated sites with the same underlying sequence motifs in all probed regions detected by AcIm and NAIN3, showing the stronger reactivity at the nucleotide immediately upstream of methylated adenine prevailing in all detected transcripts (regions with and without significant difference). Nucleotide position 0 is the m$^6$A / A site. (b) Scatter plots of R-scores for m$^6$A methylated versus non-methylated sites with the same underlying sequence motifs corresponding to (a).

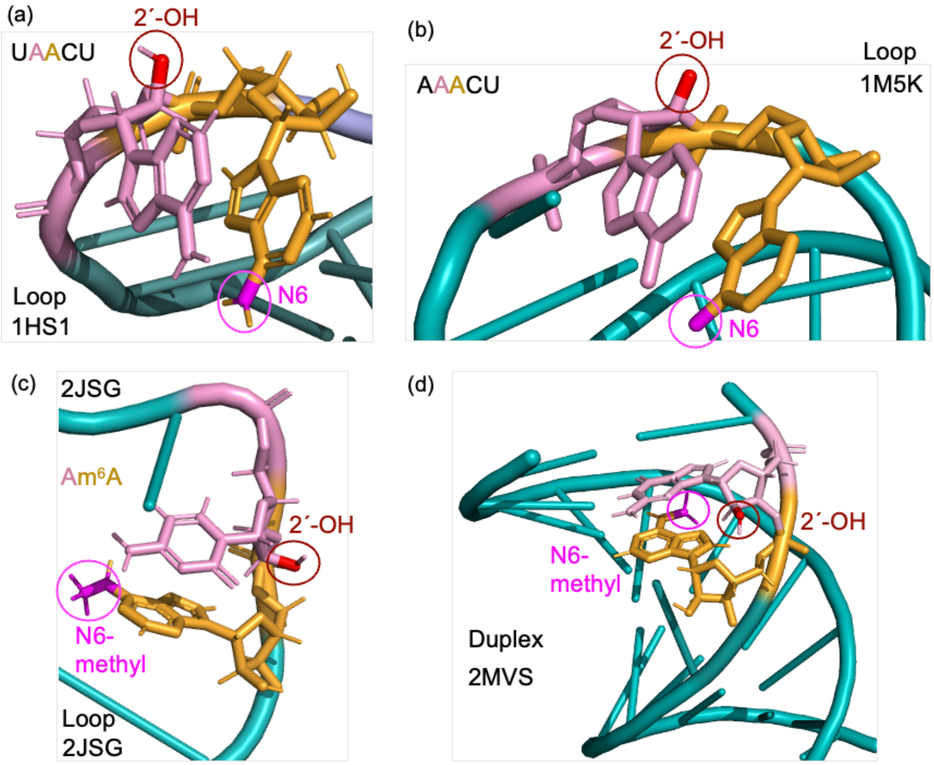


**Figure S12.** Crystal structure of RNAs with (a-b) and without (c-d) m6A in loop (a-c) and duplex (d) structures. Magenta color: N6 amine or N6-methyl group in A; red color marks the 2´-OH group of the adjacent upstream nucleotide. Methylation may enhance the stacking (2) with the adjacent upstream nucleotide to stabilize the exposure status of nearby 2´-OH group for increased probe accessibility. PDB: 1HS1, 1M5K, 2JSG, 2MVS.

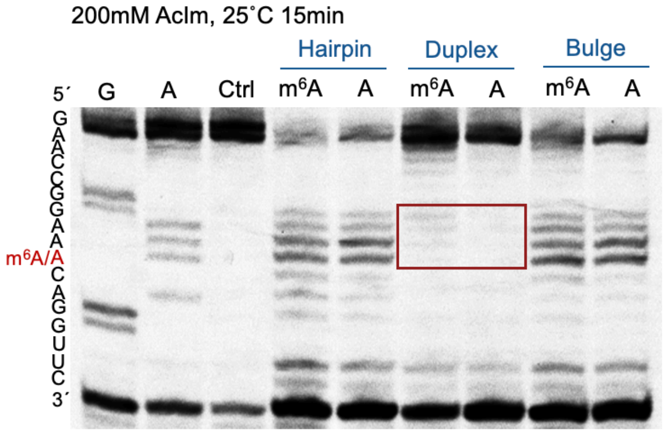


**Figure S13.** In contrast to loop regions (see main text), acylation patterns for m6A in duplexes shows little or no difference +/- methyl group. Gel analysis of reverse transcriptase (RT) stops for designed RNA structures reacting with high-concentration (200mM) AcIm at 25˚C for 15min.

Compared to Fig. 5g (main text) using 100 mM reagent, we observe that differences in acylation +/- methylation are lost at high reagent concentrations. Note that RT stops (and corresponding bands) occur at the nucleotide immediately 3' to the reacted position.

### Experimental Procedures

**Activation of acylimidazole probes.** The acylimidazole probes were generated immediately before the profiling experiments. NAIN3 was synthesized according to previously reported protocols and prepared as a 2M stock (1:1 with imidazole) in DMSO. $^1$H-NMR (400 MHz, d6-DMSO) δ=8.81 (dd, $J$ = 4.9, 1.7 Hz, 1H), 8.13-8.08 (m, 2H), 7.62 (t, $J$ = 1.5 Hz, 1H), 7.56 (dd, $J$ = 7.8, 4.9 Hz, 1H), 7.13 (dd, $J$ = 1.7, 0.8 Hz, 1H), 4.58 (s, 2H). AcIm and AcIN3 were synthesized according to the protocols below. Briefly, the carboxylic acid precursor (1.0 equiv.) was dissolved in dry DMSO as a 2 M solution. To it, 1.3 equivalents of 1,1'-carbonyldiimidazole (CDI) powder was added at room temperature and mixed until bubbling ceased. The resulting solution was stored and used as a 2 M acylimidazole stock without further purification. AcIm: 1H-NMR (500 MHz, d6-DMSO) δ=8.40 (s, 1H), 7.69 (d, J = 1.5 Hz, 1H), 7.07 (d, J = 1.6 Hz, 1H), 2.62 (s, 3H).

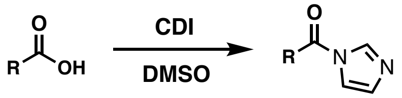


**Note:** although acetylimidazole is commercially available, it hydrolyzes in contact with moisture in the air, and thus can become less active and somewhat acidic over time as a bottle is opened repeatedly. Therefore, we recommend freshly preparing AcIm from $Na_2SO_4$-dried acetic acid and anhydrous DMSO using the above procedure for the current application.

**Cell culture, cell harvesting, probe treatment, and total cellular RNA isolation.** HEK293 cells were grown in DMEM media containing 10% FBS on 15-cm plates until they reached ~90% confluency. Cells were first harvested by washing with 10 mL of warm 1x PBS and gently scraped with 4 mL of 1x PBS. ~2x10$^6$ cells were transferred to a 15-mL falcon tube. The supernatant was removed by centrifugation at 1000*g* for 2 minutes at 25˚C. Cell pellets were resuspended in 450μL 1xPBS solution and 50μL 10x probing reagents (1M NAIN3, 1M AcIm, 2M AcIN3, and DMSO) were added. The mixtures were incubated at 37˚C with rotation for 10min. Treated cells were then lysed with 6 mL of Trizol LS by vortexing for total RNA extraction. 1.2 mL of chloroform was then added. The resulting mixture was vortexed and incubated at room temperature for 5 min, followed by centrifugation at 2500g for 15 min at 4˚C. The aqueous phase was then mixed with 1× volume of 96% ethanol and purified using a Quick-RNA MidiPrep kit (Zymo).

**Isolation of ribosomal RNAs and poly(A)+ RNA fraction.** (1) 18S and 28S rRNAs were isolated with agarose gels. Total cellular RNAs mixed with 6x blue/orange loading dye (Promega) were loaded onto a 1% agarose gel followed by gel excision and purification with QIAquick Gel Extraction Kit (Qiagen). (2) poly(A)+ RNA fraction was isolated with Poly(A)Purist MAG kit (Thermo Fisher). Briefly, a 250-μg aliquot of total cellular RNA was diluted to a final volume of 400 μL with RNA storage solution to reach a total concentration of 600 μg/mL. Poly(A)+ RNAs were then purified with Poly(A)Purist MAG kit according to the manufacturer's protocol. To further desalt and clean up the RNA, eluted RNA was first lyophilized and then dissolved in 100 μL of water to proceed with purification using RNeasy Mini columns (Qiagen) according to the manufacturer's protocol. Finally, RNA was eluted twice with 50 μL of water (100 μL final volume) and stored as 500-ng aliquots at -80℃.

**Sequencing library preparation.** To prepare the library for deep sequencing, we followed the method as previously reported, including biotin conjugation of probe-labeled RNAs, RNA fragmentation, RNA 3'-end repair and ligation, cDNA synthesis via reverse transcription with barcode-encoded primers, biotin RNA-cDNA complex enrichment, cDNA purification and circularization, library amplification. For detailed experimental procedures see reference (3).

**Bioinformatics and data analysis.** We utilized our reported RBRP bioinformatics pipeline (3) based on icSHAPE (4) to calculate averaged frequency of reverse transcription stops as the R-score to represent RISP signal at each nucleotide. Briefly, the raw sequencing data was processed by demultiplexing according to the barcode, collapsing to remove PCR duplicates, primer and linker trimming. RNA library index files were built, and trimmed reads were aligned to rRNA sequences or human transcriptome. The averaged RT-stop frequency was calculated following the scripts in the pipeline. We defined the R-score as the subtraction of background reverse transcription stops (mock DMSO libraries) adjusted by the background base density from reverse transcription stops of the acylated libraries adjusted by the sample base density:

$$\text{R-score} = (\text{RT-Stop}_{sample} / \text{base\_density}_{sample}) - \alpha\ (\text{RT-Stop}_{DMSO} / \text{base\_density}_{DMSO})$$

As the treated and mock-treated libraries are sequenced differently, this parameter does not equal 1, and it was trained on human rRNA structures and set to 0.5 to maximize the correlation of R-score determined by deep sequencing and reactivity score measured in manual gel-shift experiments based on band intensity (Fig. S1). Different from original RBRP pipeline, we normalized the top 1% R-score of each reagent to 1 and bottom 1% to 0, with R-score scaling linearly in between to compare the reagents (AcIm, AcIN3, and NAIN3) among RNAs.

We fine-tuned the informatics parameters to ensure a strong correlation (Pearson correlation $r \geq 0.85$) between the R-score and manual structure-probing gels of partial 18S rRNA (Fig. S2a-b). After calculating the R-score for 18S and 28S rRNAs using the pipeline (Fig. S3), we established specific criteria to facilitate our analysis: (1) Nucleotides with an R-score greater than 0.1 were defined as efficient signals for each profile (Fig. S2c). (2) We identified nucleotides showing significant difference between AcIm and NAIN3 in gels had an absolute differential R-score larger than 0.2 in RISP profiles, which were defined as AcIm or NAIN3 dominant nucleotides for further analysis in the whole sequencing profiles. (Fig. S2d).

To calculate the distance of 2′-OH group to nearby proteins, we employed a PyMOL script called distancetoatom.py. We analyzed the closest distance of 2′-OH group in RNA to the heavy atom of proximal proteins. To calculate RNA-protein interface, we utilized another PyMOL script called interfaceResidue.py. We calculated interfaces and non-interfaces residues between RNAs and proteins based on the crystal structure of ribosome. The calculation of interfaces took the difference between the surface area of RNA-protein complex and each object and listed the located residues for further analysis. In addition, we took advantage of a web-based tool called MOLEonline for analyzing tunnels within the human ribosome (5). The $m^6A$ database that we used was from DirectRMDB that published recently (6). Bioinformatics tools such as bedtools, seqkit were employed during analysis. The molecular volume of NAIN3 and AcIm was calculated by web-based software called molinspiration and the atom-to-atom distance was calculated by software called Avogadro.

**Acylation of synthetic RNAs in the presence or absence of $m^6A$.** 100 pmole of RNA with or without 200 pmole of corresponding helper RNA for structure construction were heated in folding buffer containing 50 mM NaCl to 95℃ for 2min and step-cooled to 25℃ at a rate of 0.5℃/s to form a defined RNA structure (hairpin, duplex, bulge). To the annealed solution was added 2.4 µL 4xMOPS buffer and 1 µL 10x acylating reagents (AcIm, AcIN3, NAIN3) or DMSO to final 10µL mixture. The reaction was incubated at 25℃ for 15min and quenched by DTT, followed by ethanol precipitation of the RNA samples. The RNAs of duplex and bulge samples recovered from precipitation contained both reacted RNAs and helper RNAs. To isolate the reacted RNAs for reverse transcriptase (RT) stops analysis, we conducted a purification via 15% urea denaturing PAGE gel. The corresponding gel bands of reacted RNAs were cut out and chopped into small pieces. The gel pieces were soaked in RNase-free water and shaken on a shaker overnight in a cold room. The extracted RNA was then ethanol precipitated by adding 1/10 volume of 3M sodium acetate, pH 5.0, 0.5 µL 20µg/µL glycogen and triple the volume of cold ethanol. The purified RNAs were further analyzed by reverse transcription for acylation analysis.

**PAGE analysis of reverse transcriptase (RT) stops.** 10 pmol RNA was mixed with 6.5 pmol FAM-labeled RT Primer and 0.25 µL 10 mM dNTP mix (for sequencing lane, ddNTP:dNTP=8:1), and incubated for 5 min at 75℃, then immediately chilled on ice for 2 min. 2 µL 5x First-Strand Buffer, 1 µL 0.1 M DTT, 0.25 µL RiboLock and 0.25 µL SuperScript III (200 U/µl) were added to the final volume of 10 µL. The reaction was incubated with the following program: 25℃ for 10 min, 42℃ for 30 min, and 52℃ for 50 min. After the reverse transcription reaction, 0.5 µL 1M NaOH was added and incubated at 95℃ for 3 min to remove the RNAs. Then 10 µL loading dye (8 M Urea, 0.05% Orange G, 0.05% Bromophenol blue) was added. The mixture was denatured at 95℃ for 3 min and loaded on a denaturing 15% polyacrylamide gel. Products were separated in a gel in 1x TBE (pH 8.3, Sigma Aldrich), 20mA, ~1.5 h. The cDNA gel was visualized by fluorescence imager (iBright FL1500).
